# Combined Auditory, Tactile, and Visual fMRI Reveals Sensory-Biased and Supramodal Working Memory Regions in Human Frontal Cortex

**DOI:** 10.1101/2025.06.21.660846

**Authors:** Sean M. Tobyne, James A. Brissenden, Abigail L. Noyce, David C. Somers

## Abstract

Selectivity for sensory modality characterizes distinct subregions of the human brain, well beyond the primary sensory cortices. We previously identified frontal and posterior cortical regions that are preferentially recruited for visual vs. auditory attention and working memory (WM). Here, we extend our approach to include tactile cognition and to characterize cortical regions recruited by WM in each of three sensory modalities. The joint organization of visual-selective, auditory-selective, tactile-selective, and supramodal WM recruitment within individual subjects has not been fully investigated previously. Male and female human subjects participated in a blocked fMRI task requiring them to perform *N-*back WM judgements in auditory, visual, or tactile (haptic) modalities. We confirmed our prior reports of multiple visual-biased and auditory-biased frontal lobe regions. We also observed several bilateral tactile-selective regions abutting previously described visual- and auditory-selective regions, including dorsal and ventral precentral sulcus, the postcentral sulcus, and the anterior intraparietal sulcus. Several cortical regions were recruited by WM in all three sensory modalities in individual subjects, including precentral sulcus, inferior frontal sulcus, intraparietal sulcus, anterior insula and pre-supplementary motor area. Supramodal regions exhibited substantial overlap with visual-biased regions in frontal and parietal cortex and comparatively little overlap with tactile- or auditory-biased regions. Lastly, resting-state analyses revealed that auditory-, visual- and tactile-selective WM regions segregate into modality-specific networks that span frontal and posterior cortex. Together, these results shed light on the functional organization of sensory-selective and supramodal regions supporting higher-order cognition.

**SIGNIFICANCE STATEMENT:** Using within-subject fMRI analyses of three different sensory modalities, we identify the fine-scale architecture of sensory-biased and supramodal working memory structures within human frontal cortex. Sixteen bilateral frontal lobe regions are identified: five auditory-biased, four visual-biased, two tactile-biased, and five multiple-demand / supramodal regions. These parcels are largely distinct with the notable exception that the visual-biased regions each partially overlap with supramodal regions, suggesting a tight link between visual and supramodal working memory representations. Sensory-biased frontal lobe regions form modality-specific networks with traditionally-identified posterior sensory cortical regions. The findings demonstrate a pervasive role for sensory modality in the functional organization of frontal cortex and offer a fine-scale parcellation of the human frontal lobe regions supporting working memory.

## INTRODUCTION

The neural substrates of working memory (WM) have long been identified as a prefrontal or a frontoparietal cortical network (e.g., Pribram, et al., 1964; Courtney et al., 1997; Postle et al., 1999). Sensory WM also recruits regions specific to that sensory modality, including primary sensory cortical areas (Pasternak & Greenlee, 2005; Christophel et al., 2017). Commonly, the fronto-parietal circuitry is viewed as domain-general or supramodal, while the posterior sensory cortical areas are viewed as sensory modality-specific (Duncan, 2010; Assem et al., 2020; Rizza et al., 2024). However, the localization of WM in the human brain is largely known via functional MRI (fMRI) using group-averaging approaches that spatially blur brain activations. Therefore, many regions described as supramodal could reflect a blurring together of neighboring regions with distinct sensory modality preferences. Detailed analyses of individual subjects can reveal finer-scale functional brain organization than is observed in group-averaged studies (Michalka et al., 2015; Braga & Buckner, 2017; Gordon et al., 2017; Gratton et al., 2020; Somers et al., 2021; Kwon et al., 2025); this approach is particularly relevant in frontal cortex where functional domains are small and exhibit anatomical variations across individuals. Here, we employ vertex-level, within-subject analyses to examine the fine-scale functional organization of human frontal lobes in the support of visual, auditory, tactile, and supramodal sensory WM.

Within lateral frontal cortex our prior research has described multiple auditory-biased regions and visual-biased regions that are recruited during sensory WM tasks and that form modality-specific networks with posterior sensory cortical regions (Michalka et al., 2015; Noyce et al., 2017, 2022; Tobyne et al., 2017, 2018; Somers et al., 2021). Group-averaged analyses based on predefined regions of interest (ROIs) confirmed the existence of these sensory-biased frontal regions and placed them posterior to several key multiple-demand regions in lateral prefrontal cortex (Assem et al., 2022). In contrast, we have observed that visual-biased, but not auditory-biased frontal regions exhibit multiple-demand WM properties (Noyce et al., 2017, 2022). Here, we perform detailed within-subject analyses to better understand the role of sensory modality in the functional organization of the human frontal lobes.

The present work also incorporates tactile (haptic) WM with visual and auditory WM to map WM circuitry in each sensory modality. Prior tactile WM studies revealed activations in superior parietal lobule, inferior frontal gyrus, middle frontal gyrus, anterior insula, premotor cortex, and medial frontal cortex (Preuschhof et al., 2006; Ricciardi et al., 2006; Sörös et al., 2007; Savini et al., 2012; Kaas et al., 2013; Schmidt et al., 2017, 2021; Schmidt & Blankenburg, 2018). Prior investigations of WM that have examined two sensory modalities at a time describe frontal cortical regions as supramodal, supporting the multiple-demand account of frontal cortex in WM (Ricciardi et al., 2006; Rizza et al., 2024; Wu et al., 2018). Here, we report two bilateral tactile-biased, four bilateral visual-biased, and five bilateral auditory-biased WM regions within frontal cortex. We also observe more posterior cortical regions that preferentially support WM in the tactile modality and others that support visual or auditory WM. Frontal tactile-biased regions form a resting-state network with posterior tactile cortical regions, similar to our prior findings for visual-biased and auditory-biased networks (Michalka et al., 2015; Tobyne et al., 2017; Noyce et al., 2022). We also identify seven regions, five frontal and two parietal, that support WM across all modalities. All visual-biased frontal regions partially overlap with supramodal WM regions, but none of the auditory-biased and only one tactile-biased region exhibit this overlap. This suggests a tight relationship between visual WM processing and supramodal WM. We summarize our findings by placing all sixteen bilateral frontal lobe domains – two tactile-biased, four visual-biased, five auditory-biased, and five supramodal– on a schematic representation.

## MATERIALS AND METHODS

### Subject Recruitment

Ten subjects participated in this study (mean age 28.1 ± 5.07 years, range 22 – 35, 5 females). Subjects were recruited from Boston University and the Greater Boston area. All subjects were healthy, right-handed and had normal or corrected-to-normal vision. Subjects were screened for MRI contraindications prior to scanning. The Institutional Review Board of Boston University approved all experimental procedures. All subjects provided written informed consent in accordance with the guidelines set by Boston University.

### Stimuli and Experimental Design

Stimuli display and task timing control for the three-modality working memory (TMWM) task was performed using the Psychophysics Toolbox (v. 3.0.14; www.psychtoolbox.org; Brainard, 1997; Pelli, 1997) in MATLAB (v. 2018a; www.mathworks.org; The MathWorks, Natick, MA), and displayed using a liquid crystal display projector that back-projected onto a screen placed within the scanner bore. Figure 1a depicts the TMWM task paradigm. A white central fixation cross was presented on a black background during all task conditions. Auditory and visual stimuli matched that of Noyce et al., 2017. Visual stimuli consisted of black and white male and female faces. Faces were presented centrally and spanned ∼6.4° of visual angle. Auditory stimuli consisted of natural cat and dog vocalizations. Auditory stimuli lasted 300-600 ms and were presented through MRI-safe in-ear earphones (Sensimetrics Model S14; Sensimetrics Co., Gloucester, MA; www.sens.com). The visual and auditory stimulus sets were selected as they are not easily identified by a semantic label or name; all stimuli presented in a block are different exemplars of a subordinate category (e.g., male faces; cat sounds). The rationale for this choice is to force subjects to rely on sensory rather than semantic working memory. Tactile stimuli were administered using a custom-designed tactile stimulator used in previously published work (Figure 1b; Merabet et al., 2007). The stimulator consists of two PVC pipe sections, one nested inside the other, to create inner and outer shells. The tactile stimulator was placed on the scanner bed next to the subject where they could comfortably rest their hand on the end of the stimulator. Tactile stimuli consisted of 3D printed non-informative Braille-like raised dot patterns of increasing density (Figure 1c). Subjects felt the tactile stimuli by sweeping their right index finger forward and backward one time over a given trial’s raised dot pattern through an aperture cut into the outer shell. Although the tactile stimuli are less complex than the visual and auditory stimuli, the use of low-level stimuli pushes participants again to rely on sensory rather than semantic WM.

**Figure 1:**
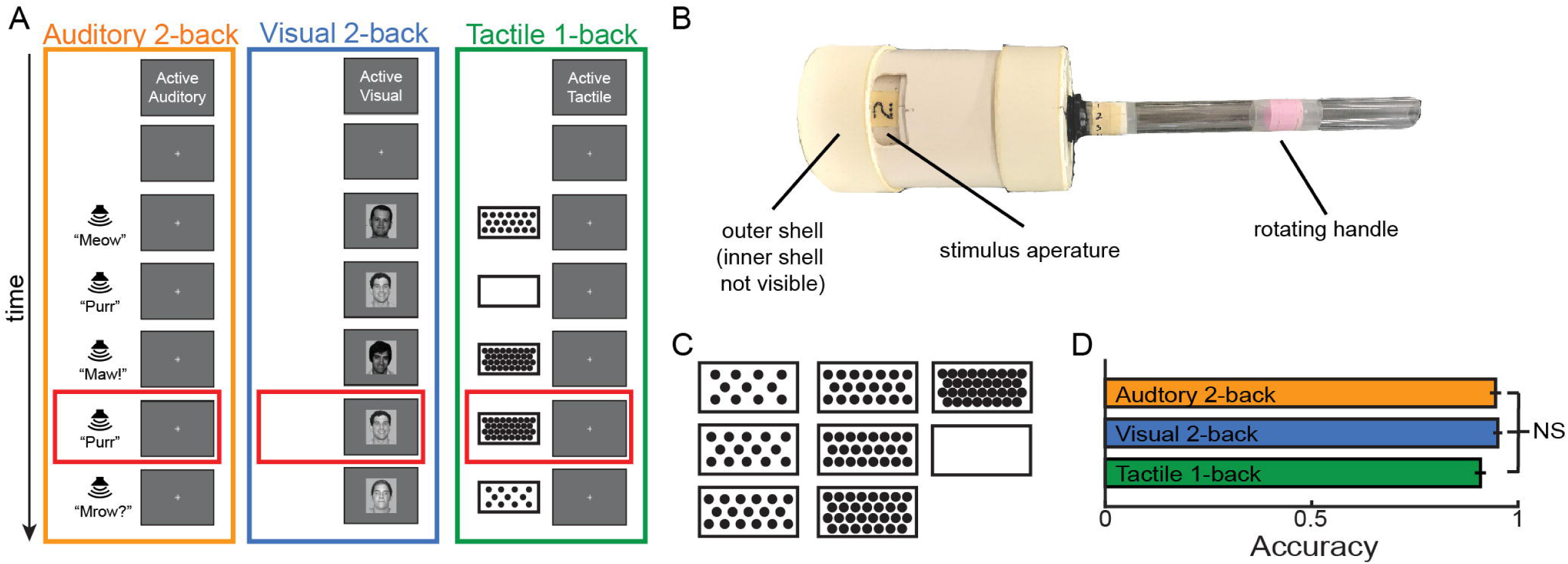
Three-Modality Working Memory Task. (a) Task protocol, (b) tactile stimulator design and (c) tactile stimuli for the TMWM fMRI task are depicted. Red boxes in (a) indicate an N-back target trial. (d) Behavioral performance during the TMWM task. NS = not significant.

An experimenter stood in the scanner room and presented stimuli to the subject by using the handle to rotate the inner shell to the appropriate stimulus. The stimulus to be presented on a given trial was converted from a numerical representation in MATLAB to computerized speech using the build in ‘talk’ command line function on a MacBook Air and conveyed to the experimenter through pneumatic headphones provided by the MRI manufacturer. The tactile stimulator was left in place for the entirety of the MRI scan session and subjects indicated responses using their left hand to make button presses on an MRI-safe response box (fORP 2-button, Cambridge Research Systems, Ltd., www.crsltd.com).

Each subject participated in 10 runs during which they performed N-back working memory tasks. Individual runs consisted of six blocks: Three active working memory blocks (1 each of auditory, tactile and visual modalities) and three passive sensorimotor control blocks (1 matched to each sensory modality). Block order was counter-balanced across runs and subjects. Blocks were 36 sec in length and were preceded by a 4 sec fixation cross presentation and then a 2 sec cue for the upcoming block; a final 4 sec fixation cross presentation was included at the end of the run. The time between stimulus onsets for auditory and visual blocks was 1200 ms in length and 30 total stimuli were presented in each block. The time between tactile presentations was 2400 ms in length and 15 total stimuli were presented. This difference in trial structure was necessitated by the latency inherent to manually operating the tactile stimulator. To roughly balance the ratio of N-back targets to total stimulus presentations, auditory and visual active blocks contained 4 N-back targets while tactile active blocks contained 3 N-back targets. Subjects performed a 2-back working memory judgement in auditory and visual active blocks but a 1-back judgment on tactile active blocks because behavioral piloting revealed that a tactile 2-back was more difficult than an auditory or visual 2-back. Passive blocks were presented in the same manner as active blocks except that they contained no N-back repeats. During tactile passive blocks, subjects were presented only with the blank tactile stimulus (see Figure 1C) but were instructed to make the same sweeping motion across the stimulus with their index finger. This instruction was given to balance the somatomotor neural response across tactile task conditions.

### Magnetic Resonance Image Acquisition

MRI data were acquired using a Siemens Prisma 3-Tesla MRI scanner located at the Cognitive Neuroimaging Center at Boston University using a 64-channel head coil. High-resolution T1-weighted (T1w) multi-echo MPRAGE (repetition Time (TR) = 2780 ms; echo time (TE) = [1.32, 3.19, 5.11, 7.03] ms; flip angle (FA) = 7°; 0.8 mm isotropic voxels; 224 slices; field of view (FOV) = 256 mm x 256 mm x 180 mm; in-plane GRAPPA (R) = 2; bandwidth (BW) = 650; acquisition time (TA) = 8 min 7 sec) and T2-weighted (T2w; TR = 3200 ms; TE = 564 ms; FA = variable; 0.8 mm isotropic voxels; 224 slices; FOV = 256 x 256 x 180 mm; R = 2; BW = 651; TA = 6 min 46 sec) bandwidth-matched structural images were acquired. All functional data were acquired using a blipped-CAIPI simultaneous multi-slice sequences (SMS) gradient-echo echo-planar imaging (EPI) pulse sequence (Moeller et al., 2010; Feinberg et al., 2010; Setsompop et al., 2012; Xu et al., 2013) sensitive to blood oxygen level dependent contrast. The ten task fMRI acquisitions (TR = 2000 ms; TE = 35 ms; FA = 80°, 2.2 mm isotropic voxels; 69 slices; FOV = 207 x 207 x 152 mm; SMS = 3; R = 0; 128 TRs; TA = 4 min 16 sec) and three resting-state acquisitions (TR = 650 ms ; TE = 34.8 ms; FA = 52°; 2.3 mm isotropic voxels; 64 slices; FOV = 207 mm x 207 mm x 148 mm; SMS = 8; R = 0; 554 TRs; TA = 6 min 10 sec) were acquired. Pairs of protocol-matched reversed phase encoding (i.e. anterior-to-posterior and posterior-to-anterior) spin echo field maps were also acquired for subsequent EPI de-warping. During resting-state acquisition, subjects were instructed to fixate on a centrally presented crosshair, remain as still as possible and allow their minds to wander.

### MRI Preprocessing

Preprocessing of all MRI data was accomplished using modified versions of the Human Connectome Project (HCP) Minimal Preprocessing Pipeline (Glasser et al., 2013; MPP; v. 3.20; https://github.com/Washington-University/HCPpipelines). Briefly, the high resolution T1w and T2w images were corrected for gradient nonlinearity distortion, ACPC aligned, skull-stripped, corrected for readout distortion, aligned in native T1w space and bias field corrected. The final transformation was accomplished in a single interpolation step by concatenating the transformations from each preprocessing stage. The preprocessed T1w and T2w images were then processed with FreeSurfer (HCP-modified v. 5.3; https://surfer.nmr.mgh.harvard.edu/pub/dist/freesurfer/5.3.0-HCP/) to construct a native surface mesh representing the cortical sheet. Note that HCP’s MPP additionally transforms all data to MNI152 template space, however we performed all individual subject analyses in native space. A final preprocessing step unified data into GIFTI surfaces and CIFTI scalar data and calculated estimated myelin content using the T1w and T2w images (Glasser & Van Essen, 2011; Glasser et al., 2014).

Task and resting-state data each followed similar preprocessing workflows, again using the HCP MPP. Briefly, both were corrected for gradient nonlinearity distortion, subject head motion, and EPI distortion using the spin echo field maps before registration to native T1w space in a single spline interpolation step. Functional data was then resampled to the individual’s reconstructed cortical surface, using the “partial volume weighted ribbon-constrained” resampling algorithm and noisy voxel exclusion (Glasser et al., 2013). Upon resampling to the surface, task and resting-state data were spatially smoothing along the geodesic surface with a 2 mm FWHM Gaussian kernel. Task fMRI data were then analyzed with FreeSurfer’s FS-FAST software package (v. 5; https://surfer.nmr.mgh.harvard.edu/fswiki/FsFast).

Resting-state fMRI data were separately postprocessed with in-house developed MATLAB scripts to accomplish linear interpolation across high-motion time points (> 0.6 mm framewise displacement; Power et al., 2012, 2014; Carp, 2013; Hallquist et al., 2013), application of a fourth-order Butterworth temporal bandpass filter (0.009 < *f* < 0.08 Hz), temporal denoising by multiple regression of the mean grayordinate signal and 24 motion confound regressors (Friston et al., 1996), and high-motion timepoint censoring via deletion. The three resting-state acquisitions for each subject were then temporally demeaned and concatenated to create a single dense timeseries.

### TMWM fMRI Task Analysis

Task fMRI data were analyzed with standard FS-FAST procedures. Fully processed task fMRI data were converted from CIFTI format to GIFTI format for use with FS-FAST. A general linear model (GLM) was fit to each vertex with regressors that matched the task conditions and orthogonalized confound regressors derived from a singular value deconstruction of the motion parameters calculated during motion correction. Cue timepoints were included as nuisance regressors to control for task reorienting effects. A canonical hemodynamic response function modeled by a gamma function with a delay of δ = 2.25 and decay time τ = 1.25 (Boynton et al., 1996) was convolved with each regressor prior to GLM fitting. Conditions of interest were then contrasted against one another to derive *t* statistics and associated *p* values for each vertex. As in prior studies (Michalka et al., 2015; Noyce et al., 2017, 2022), active WM task conditions for each sensory modality were contrasted against one another to localize regions that were preferentially recruited for one modality over another. In previous work only auditory and visual modalities were used and thus could be directly contrasted, here, however, we conducted three separate contrasts representing the three different sensory modalities (auditory > visual and tactile; tactile > visual and auditory; visual > auditory and tactile). We also separately performed contrasts for WM > passive sensorimotor control for each modality, as a secondary control to screen out regions without a specific WM component. The final *z*, *t,* and *p* statistic maps were converted to CIFTI format for visualization with Connectome Workbench.

#### Assessment of Sensory Modality-biased Responses

The specification of the contrast of the auditory, tactile and visual active conditions dictates that significant results can be achieved by one modality surpassing only one of the two other modalities – allowing for the possibility of overlapping significant results across analyses. To assess sensory modality-biased recruitment, a disjunction analysis was conducted to localize uniquely recruited cortex for each sensory modality. Using Connectome Workbench’s (Marcus et al., 2011; CWB; https://www.humanconnectome.org/software/connectome-workbench) wb_command and the -cifti-math function, we first thresholded each individual contrast of one modality over the other two modalities at a liberal *p* = 0.05 (positive activation only) and them masked out any vertices that remained in either of the two other contrasts (e.g. all significant visual WM > auditory WM and tactile WM vertices and all significant tactile WM > auditory WM and visual WM vertices were masked from the auditory WM > tactile WM and visual WM map). The 3 resulting maps represented vertices that were significantly recruited (compared to the combination of the other two modalities) only for each modality’s active condition. These modality-biased maps were then used to hand draw regions of interest (ROIs) using CWB wb_view visualization program by defining a border that encompassed each ROI. This border was then converted to a ROI and finally a label using CWB wb_command’s -border-to-rois and - metric-label-import commands, respectively. This procedure is not able to account for ‘holes’ of sub-threshold vertices within the hand-drawn border, so masking steps were performed using custom MATLAB functions to ensure that 1) only significantly active vertices from the disjunction maps were included in hand-drawn labels and 2) no abutting labels mistakenly contained the same vertices.

The identification or labeling of specific visual-biased and auditory-biased ROIs was guided by comparing their position and modality preference with the 3 bilateral visual-biased and 5 bilateral auditory-biased frontal ROI identified in our prior studies (Michalka, 2015; Noyce et al., 2017; 2022). Although the three-way contrasts employed here tended to yield slightly smaller ROIs than obtained in the vision vs. auditory studies, the MNI coordinates of the centroids and the preferred modalities were very well matched. For the tactile-biased and supramodal ROI definitions, we were guided by the hotspots in the probabilistic ROI maps, and then again used modality preference and anatomical location to label specific regions in individuals.

Task-based ROI analyses were conducted for WM contrasts by extracting percent signal change (PSC) values from all voxels within each ROI, averaging across all 10 acquisitions. PSC was calculated relative to baseline activation levels during each modality’s matched sensorimotor control conditions. We conducted separate repeated measures ANOVAs for each ROI to test the relationship between hemisphere and modality. Following Mauchly’s tests for violations of sphericity, lower bound corrections for the degrees of freedom for the *F*-test were applied. ROI area was calculated using the CWB wb_command -surface-vertex-area command and each subjects’ native midthickness surface. If ROI-based PSC analyses revealed that an ROI was not recruited by its like-modality WM contrast it was removed from the ROI set for further analyses. This screening test was performed to eliminate ROIs that did not contribute to WM processes or could have arisen solely due to greater deactivation of one sensory modality over another in default mode regions. Note that the WM > passive sensorimotor control contrast within an ROI defined by a contrast of one modality against the other two modalities is positively biased due to both contrasts being computed as a function of the same condition. As a result, a null result for the WM contrast provides strong evidence that an ROI is not recruited by WM.

The location and size of individual ROIs, as well as the standard deviation across individuals is summarized in Table 1. The MNI coordinates of the ROI centroid were used in the location calculations. For visualization purposes, we created probabilistic ROI maps by converting binarized and thresholded (see Methods) individual-subject hemispheric activation maps from native space to the fs_LR brain surfaces. The probability of any surface vertex belonging to any of the activated regions was calculated by quantifying the percentage of subjects for which activation was present. These vertex-level histograms were mapped on the fs_LR brain in Figures 4 and 11. These probabilistic maps in turn were used to guide the labeling of tactile-biased and supramodal ROIs in individual subjects. Not surprisingly, nearby probabilistic ROIs with different modality preferences can overlap; however, in individual subjects our sensory-biased ROIs contained no overlap. To better convey the functional organization for a typical individual subject, we hand drew schematic illustrations onto the fs_LR brain (Figures 5, 12). Our definition of supramodal ROIs does not exclude the possibility of overlap with sensory-biased ROIs and Figure 12 attempts to characterize the overlaps observed.

**Table 1:**
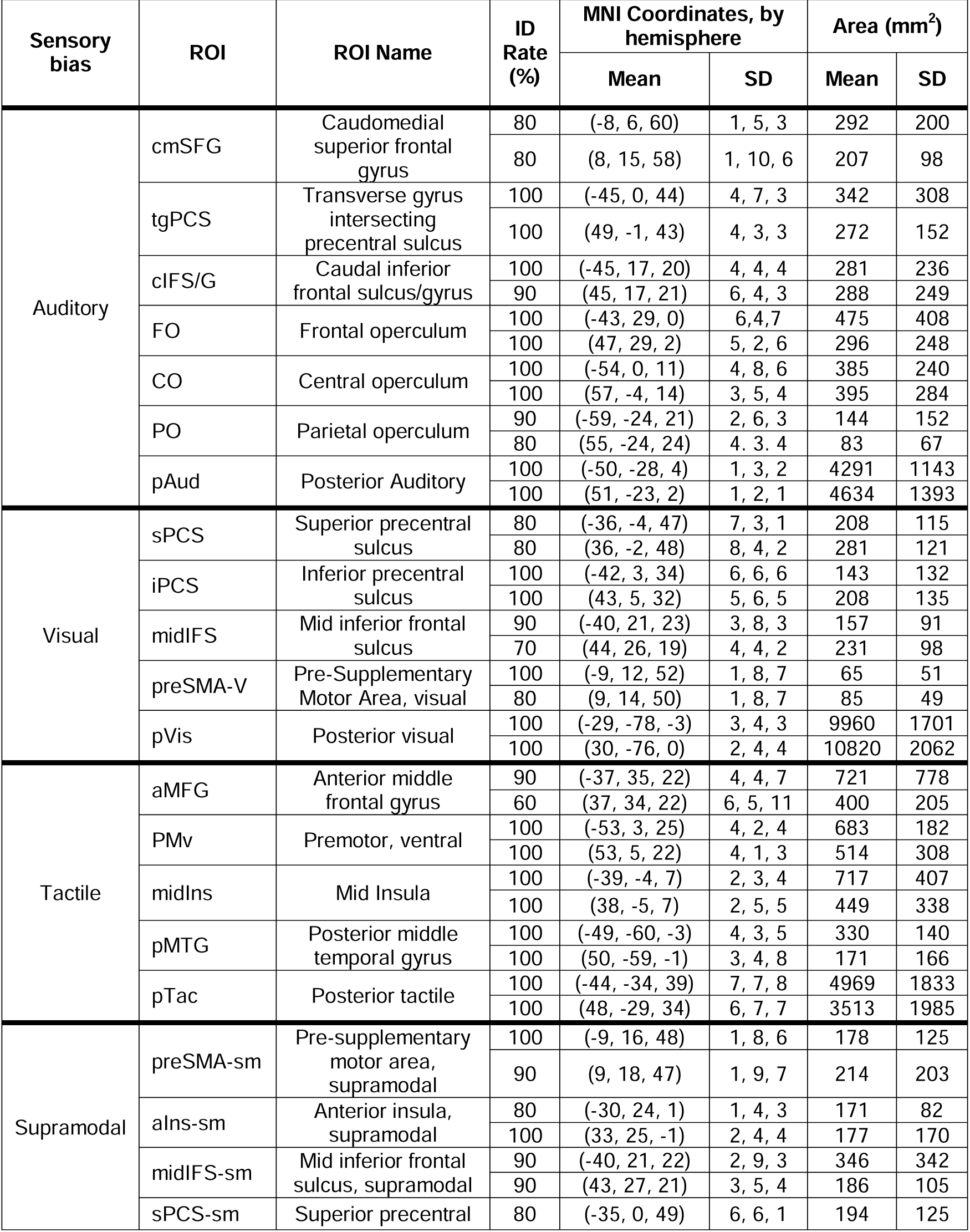
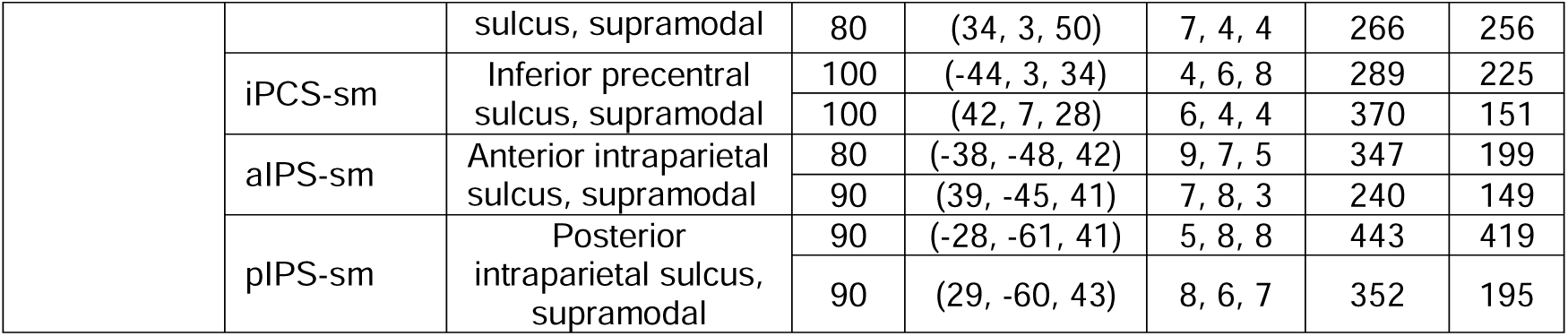
Anatomical localization of sensory-biased and supramodal ROIs. Regions of interest were defined within individual subjects in both hemispheres. The identification rate (% of subjects), MNI coordinates of the centroids, and sizes are summarized for each ROI.

#### Group Analysis of Sensory-biased WM

The standard deviation of region centroids for sensory-biased sPCS, tgPCS, iPCS and cIFS/G is nearly 90% of their geodesic width, indicating that there is substantial frontal lobe anatomical variability across individual subjects (Michalka et al., 2015; Noyce et al., 2017, 2022; Tobyne et al., 2017). To highlight the importance of single subject analyses for the expanded set of sensory-biased regions identified in this study, we also conducted group-level analyses of auditory active > visual active, as well as tactile active > auditory active and tactile active > visual active. Individual *z-*statistic maps from the FS-FAST GLM fitting procedure were converted to GIFTI surfaces in common fs_LR space, concatenated and analyzed with FSL’s Permutation of Linear Models (PALM; Winkler et al., 2016) tool (1024 sign-flips, FWE-corrected, cluster extent threshold *z* = 3.1, Bonferroni correction across hemispheres).

#### Assessment of Supramodal WM Responses

Supramodal WM activation was examined at the cortical vertex and ROI level in individual subjects by combining the working memory activation findings for each of the three sensory modalities into a 3-dimensional space (auditory, tactile, visual). Separately for each subject, vertex-wise *z*-statistics for each sensory modality WM condition > its matched passive sensorimotor control were thresholded at *p* < 0.05, mapped to 3D Cartesian space and then projected to the unit sphere to normalize overall activation differences across vertices:

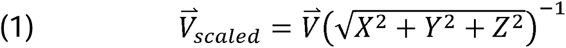

where 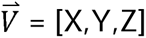 = [auditory WM *z* statistic, tactile WM *z* statistic, visual WM *z* statistic]. In this space, vertices located at or near the poles at the positive end of each axis represent uni-sensory-biased recruitment (e.g. point [1,0,0] is auditory-biased only; see Figure 8A), vertices located along arcs defined by pairs of the primary axes (i.e. X➔Y, X➔Z, Y➔Z) represent bi-sensory recruitment, while vertices located in a circle projected onto the positive octant of the sphere (highlighted sector in Figure 8A) represent supramodal recruitment. Vertices contained within this supramodal region of the sphere were mapped back to individual subject cortical surface space and separated into individual ROIs. For visualization in spherical polar space, each point was multiplied with the vector [255 0 0; 0 255 0; 0 0 255] – thus encoding each point’s degree of sensory-biased recruitment in RGB color space (R = auditory, G = tactile, B = visual).

A supramodal index (SMI), with range of −1 to 1, was computed to quantitatively assess the degree of overlapping supramodal WM recruitment across modalities for each cortical vertex. SMI was defined as the cosine of the spherical distance between each supramodal point in polar space, 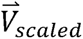, and an idealized supramodal point equidistant between auditory, tactile and visual uni-sensory-biased point, 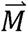, which is found by taking the inverse tangent of the Euclidean norm of the cross product between the output of Equation 1 and 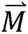 (Equation 2). SMI is normalized by the constant, *c*, such that the metric ranges from [−1,1], where −1 represents antagonistic sensory interactions, 0 represents sensory-biased working memory responses in a single modality and 1 represents a fully supramodal working memory response:

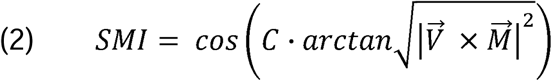

where 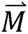 is the idealized supramodal point and *c* is the normalization constant calculated as:

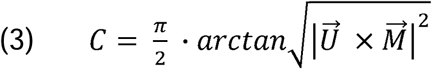

where 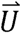 is any uni-sensory point on the sphere (i.e. [1 0 0], [0 1 0] or [0 0 1]). SMI was computed for each vertex over the entire cortical surface and then averaged for each supramodal and sensory-biased ROI.

### Resting-state Network Analysis

To test the network-level organization between identified sensory-biased ROIs seed-to-seed functional connectivity analyses were conducted. For each subject, the mean time courses extracted from each ROI, and Pearson’s correlation coefficients were calculated between all pairs of ROI time courses. This analysis was performed on the set of identified sensory-biased ROIs. Correlation values were converted to *z*-values with Fisher’s *r*-to-*z* transformation prior to statistical analysis. Group average values were converted back to *r* values for reporting purposes and visualized as group aggregate adjacency matrices. For visualization, adjacency matrices were liberally thresholded at *r* < 0.2 to remove low and likely non-significant correlation coefficients.

### Hierarchical Cluster Analysis

Using each region’s connectome fingerprint, we performed hierarchical cluster analysis (HCA) to investigate how the modality-based ROIs segregate into networks. The seed-to-seed functional connectivity between the auditory-, tactile- and visual-biased ROIs was used as the input data for clustering. Connectivity profiles were averaged across the group to create an average connectivity profile for each ROI. These group average connectivity profiles were then used as features for hierarchical cluster analysis. The Euclidean distance between each vector was calculated and Ward’s linkage algorithm, which merges pairs of clusters that minimize the total sum of squares for the node to centroid distances, was applied to generate a cluster tree. Because HCA will provide a clustering solution regardless of whether any true clusters exist, a bootstrapping procedure (Dosenbach et al., 2007; Brissenden et al., 2016; Tobyne et al., 2017; Chrastil et al., 2018) was used to validate the tree branch points. Higher values indicate great confidence in the leaves contained under each branch point. A total of 1000 bootstraps were generated by randomly sampling, with replacement, from the pool of 10 individual subject sets of connectivity matrices. The resampled matrices were averaged and clustered as above to create a set of 1000 bootstrapped cluster trees. Confidence values at each branch point of the primary HCA dendrogram were computed by calculating the proportion of bootstrap trees where the subtree clustered the same ROIs as the primary HCA.

### Statistical Analysis

All statistical analyses were performed with MATLAB (v. 2018a; the MathWorks, Native, MA). Behavioral performance was tested for differences between modalities with a one-way ANOVA and Tukey’s post-hoc tests for significance. As previously noted, percent signal change analyses within and across modalities were conducted separately for each ROI with repeated measures ANOVAs, including Mauchly’s test for sphericity and lower bound correction of the *F*-test where indicated. ROIs were combined across hemispheres where a main effect for hemisphere was not observed. N-way ANOVAs were used to test for main effects of ROI and sensory modality. A similar procedure was used to test hemispheric effects for the following analyses: proportion of sensory-biased vertices within supramodal ROIs, proportion of supramodal ROIs within sensory-biased ROIs and per-ROI SMI. Student’s *t-*tests were used to test all analyses against the null hypothesis that the mean was equal to zero. Calculated *p*-values were corrected for multiple comparisons with the Holm-Bonferroni method and considered significant if the corrected *p*-values were lower than 0.05.

## RESULTS

### Behavioral Task Performance

Working memory task performance for each modality was high across subjects (auditory: mean accuracy = 0.95 ± 0.03; visual: mean accuracy = 0.96 ± 0.02; tactile: mean accuracy = 0.92 ± 0.02). Group analysis indicated that performance was not significantly different across modalities (one-way ANOVA: *F*_2,7_ = 3.36; Tukey’s HSD post-hoc: all *p* > 0.06) (Figure 1D). Note that responses for 1 subject were not recorded due to a malfunction with the response box.

### Auditory- and Visual-biased Region of Interest Identification

To define a region as a *sensory-biased WM* region, we required that it be more strongly recruited for working memory in one sensory modality than for the other two. Therefore, the primary contrast for each sensory modality was Modality A WM vs. (Modality B WM + Modality C WM). This analysis was performed vertex-wise across the cortical hemispheres for each subject and was used to define candidate sensory-biased ROIs (Figure 2a,b). We then screened out any regions which were not more strongly activated during WM for that modality than for passive stimulation in that modality. This screening was performed at the ROI level. Figure 2a displays the auditory- and visual-biased activation maps for 3 example subjects. Note that warm colors (yellow, red) indicate auditory WM > (visual WM + tactile WM), while cool (blue, turquoise) colors indicate visual WM > (auditory WM + tactile WM). Figure 2b depicts the hand-drawn auditory- and visual-biased ROIs for these 3 subjects.

**Figure 2:**
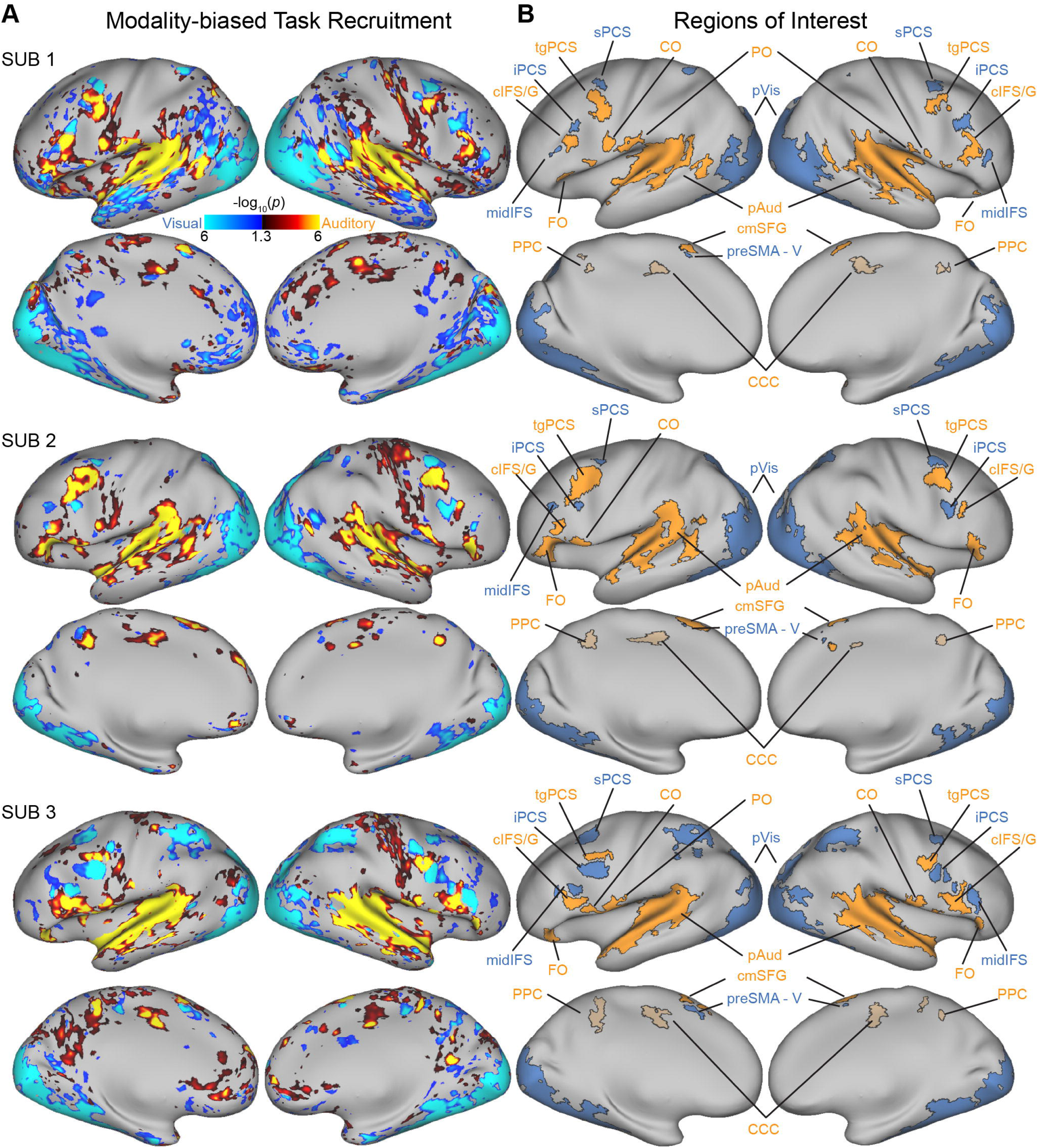
Auditory- and Visual-biased ROI Identification and Preferred Modality WM Recruitment. (a) Combined auditory-biased (hot colors) and visual-biased (cool colors) maps for 3 example subjects. Auditory-biased regions are defined by the contrast of Auditory WM > (Visual WM and Tactile WM), while visual-biased regions are defined by the contrast of Visual WM > (Auditory WM and Tactile WM) (see Materials and Methods). Both contrasts are displayed on the same figure panels (b) Final hand drawn ROIs for the same 3 example subjects.

We observed 5 auditory-biased frontal ROIs: transverse gyrus intersecting the precentral sulcus (tgPCS), caudal inferior frontal sulcus/gyrus (cIFS/G), frontal operculum (FO), central operculum (CO), and the caudomedial extent of the superior frontal gyrus (cmSFG), along with a very extensive posterior auditory complex (pAud). We also observed 3 visual-biased frontal ROIs: superior precentral sulcus (sPCS), inferior precentral sulcus (iPCS) and midway along the inferior frontal sulcus (midIFS), along with a very extensive posterior visual complex (pVis). These sensory-biased ROIs correspond in modality preference in MNI location to those reported in Noyce et al., 2022 and we apply the same nomenclature here. This also provides independent replication of these regions (see also Michalka et al., 2015; Noyce et al., 2017; Assem et al., 2022).

Three additional auditory-biased regions not addressed in our previous work were also found in most subjects, including auditory-biased parietal operculum (PO), central cingulate cortex (CCC) and precuneus/posterior cingulate (P/PC), while one small additional visual-biased region, pre-supplementary motor area visual (preSMA-V) was noted. CCC and P/PC were subsequently found to not show recruitment during auditory WM (vs. auditory sensorimotor control condition) and were eliminated form further analysis (see Figure S1). No visual ROIs failed to show recruitment during visual WM (vs. visual sensorimotor control). In summary, the visual-biased WM regions identified include bilateral sPCS, iPCS, midIFS, pre-SMA-V, and the large posterior pVis, while the auditory-biased WM regions identified include bilateral tgPCS, cIFS/G, FO, CO, cmSFG, and the posterior regions PO and pAud (See Table 1 for additional ROI details). This portion of the study confirms all three visual-biased and five auditory-biased bilateral frontal lobe regions reported in Noyce et al., (2022).

### Tactile-biased Region of Interest Identification

Analysis of the tactile-biased WM contrast (tactile WM > visual WM & auditory WM) indicated that 7 candidate tactile-biased ROIs could be reliably identified across subjects: anterior middle frontal gyrus (aMFG), premotor dorsal (PMd), premotor ventral (PMv), middle insula (midIns), posterior middle temporal gyrus (pMTG), presupplementary motor area tactile (preSMA-T), and posterior tactile (pTac). pTac is an extensive posterior tactile complex, with the name chosen to mirror pAud and pVis. Figure 3 depicts the tactile-biased disjunction activation map (panel a) and resulting hand drawn ROIs (panel b) for the three example subjects in Figure 2 – along with their auditory- and visual-biased ROIs. We also noted activity visible in the left hemisphere primary motor cortex of all subjects that likely corresponds to right hand index finger movement and right hand stimulation by the tactile stimulator. This region was drawn as the Hand Exclusion (HE) region and is described but not analyzed further. PMd and pTac are continuous with this HE region in nearly all subjects. When drawing PMd and pTac, borders were traced along the crown of the precentral gyrus and postcentral gyrus, respectively, to provide a dividing line between the putative tactile-biased cognitive region and purely somatomotor cortex. PMd and preSMA-T did not show recruitment during tactile WM (in the contrast of tactile WM vs tactile sensorimotor control) and were eliminated from further analysis (see Figure S1). All subjects’ final set of auditory, visual and tactile ROIs are displayed in Figure S2 and the MNI coordinates of their centroids and average ROI sizes are reported in Table 1. Major anatomical landmarks for these brain views are illustrated in Figure S3.

**Figure 3:**
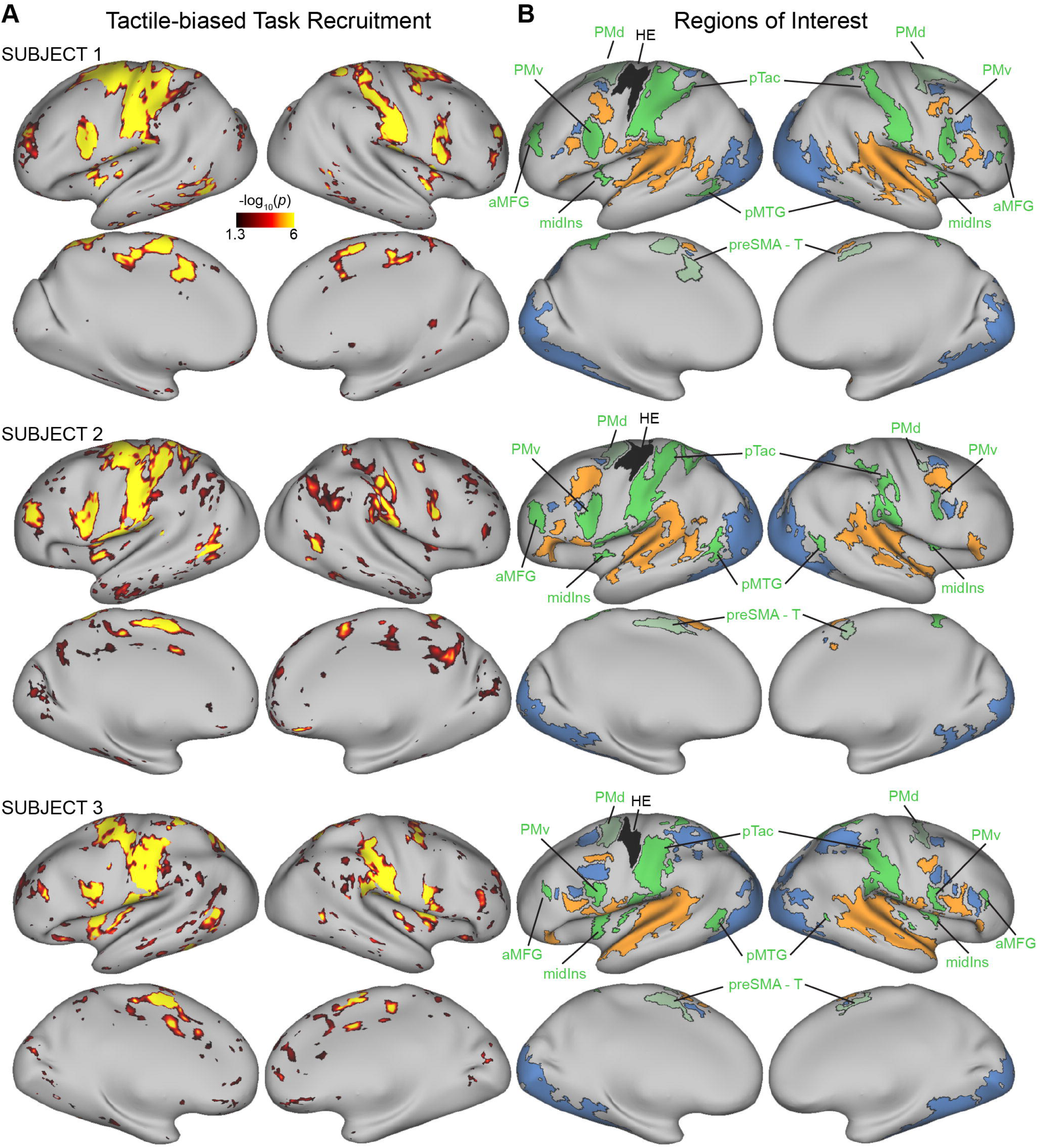
Tactile-biased ROI Identification. (A) Tactile-biased (tactile WM > visual WM & auditory WM) maps for 3 example subjects. (B) Final hand drawn ROIs for the same 3 example subjects.

Figure 4 provides a visualization of the topological layout of probabilistic auditory- and visual-biased ROIs and the probability maps of tactile-biased ROIs. PMv, midIns and pTac showed greater alignment across subjects, relative to aMFG and pMTG. Probabilistic auditory-biased tgPCS, cIFS/G, CO and pAud and probabilistic visual-biased iPCS, midIFS and pVis generally spare the regions of high tactile overlap (orange to yellow) and overlap only with tactile probabilities indicating 1-2 subject overlap. Visual-biased sPCS and the visual and auditory components on the medial surface, preSMA-V and cmSFG, showed no overlap with the tactile probabilistic maps. Only auditory-biased PO showed significant overlap with a tactile-biased probability map (pTac). This owes to the fact that pTac often surrounded or directly abutted the PO region in individual subjects. Thus, at the group aggregate level, the PO region appears to overlap with pTac.

**Figure 4:**
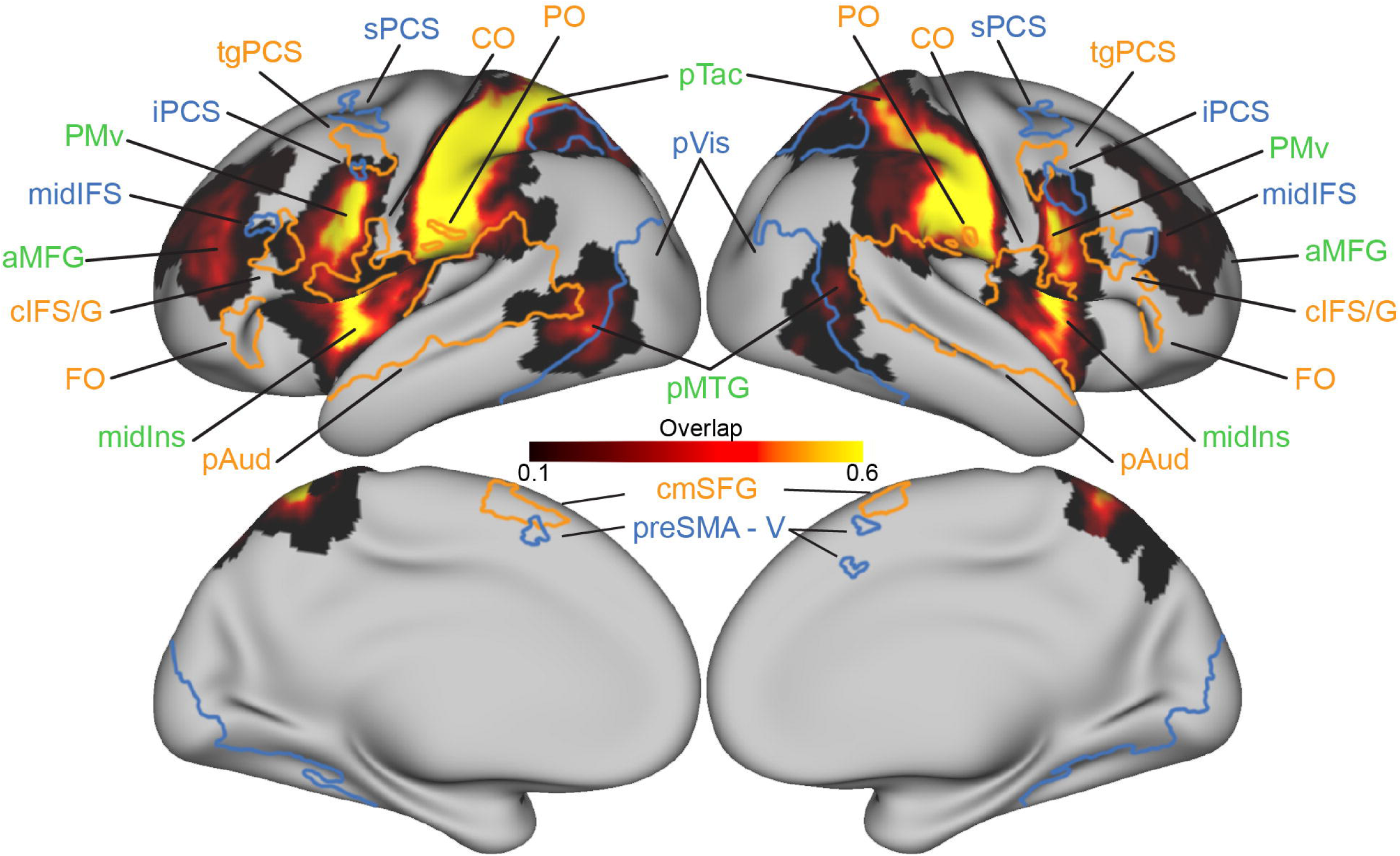
Probabilistic Tactile-biased ROI Map and Probabilistic ROIs for auditory- and visual-biased regions. The probabilistic overlap map for tactile-biased ROIs is displayed for the range of 10% (black = 1 subject) to 60% or greater subject overlap (yellow => 6 subjects). Auditory- and visual-biased ROIs created by thresholded at a 20% subject overlap are displayed in corresponding orange and blue borders.

### Schematic Layout of Cortical Sensory-biased Regions

Figure 5 presents an updated schematic layout of sensory-biased WM regions for vision, audition, and touch. This hand-drawn illustration attempts to summarize our current best understanding of the topographical layout of sensory-biased WM regions across the cortex in a typical individual subject and serves as the scaffolding with which we approached all remaining analyses.

**Figure 5:**
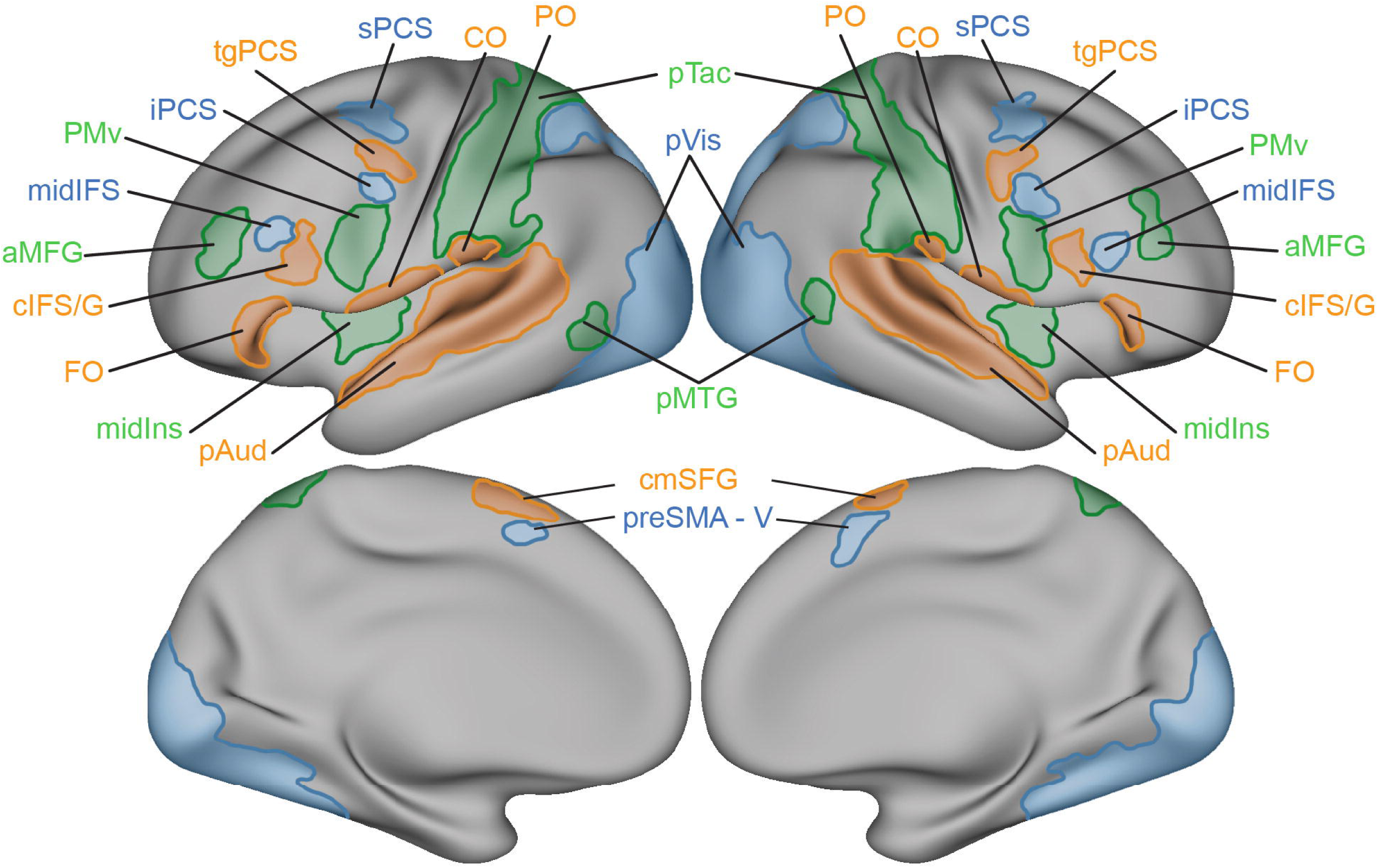
Schematic Representation of Sensory-biased ROIs. Visual-biased regions (blue), auditory-biased (orange), and tactile-biased (green) are depicted in their approximate location and size to serve as a schematic representation of the topography of sensory-biased WM regions across the cortex.

### Functional Connectivity Network Analysis

The average group adjacency matrix resulting from seed-to-seed functional connectivity revealed clear within-modality network structure, suggesting that within-modality regions form modality-specific intrinsic networks (Figure 6A). Within-modality grouping is apparent along the diagonal of the adjacency matrix for within-hemisphere auditory-, tactile- and visual-biased regions, as well as across hemisphere. Generally, the pattern of interhemispheric connections reflects strong homotopic connections, except perhaps in the auditory-biased regions, where stronger lateralization effects are observed. Auditory network lateralization is consistent with prior functional connectivity results suggesting that the auditory network makes robust connections to the language network, which exhibits strong lateralization. (Tobyne et al., 2017; Noyce et al., 2022). Notably, auditory-biased CO and PO demonstrated low correlations with frontal auditory-biased ROIs, but strong connectivity with pAud. Nearly all the tactile-biased and visual-biased ROIs possessed moderate or strong within-modality correlations. Visual-biased ROIs possessed the least cross-modality functional connectivity with auditory- or tactile-biased ROIs. We observed moderate correlations between tactile-biased midIns and auditory-biased CO and PO.

**Figure 6:**
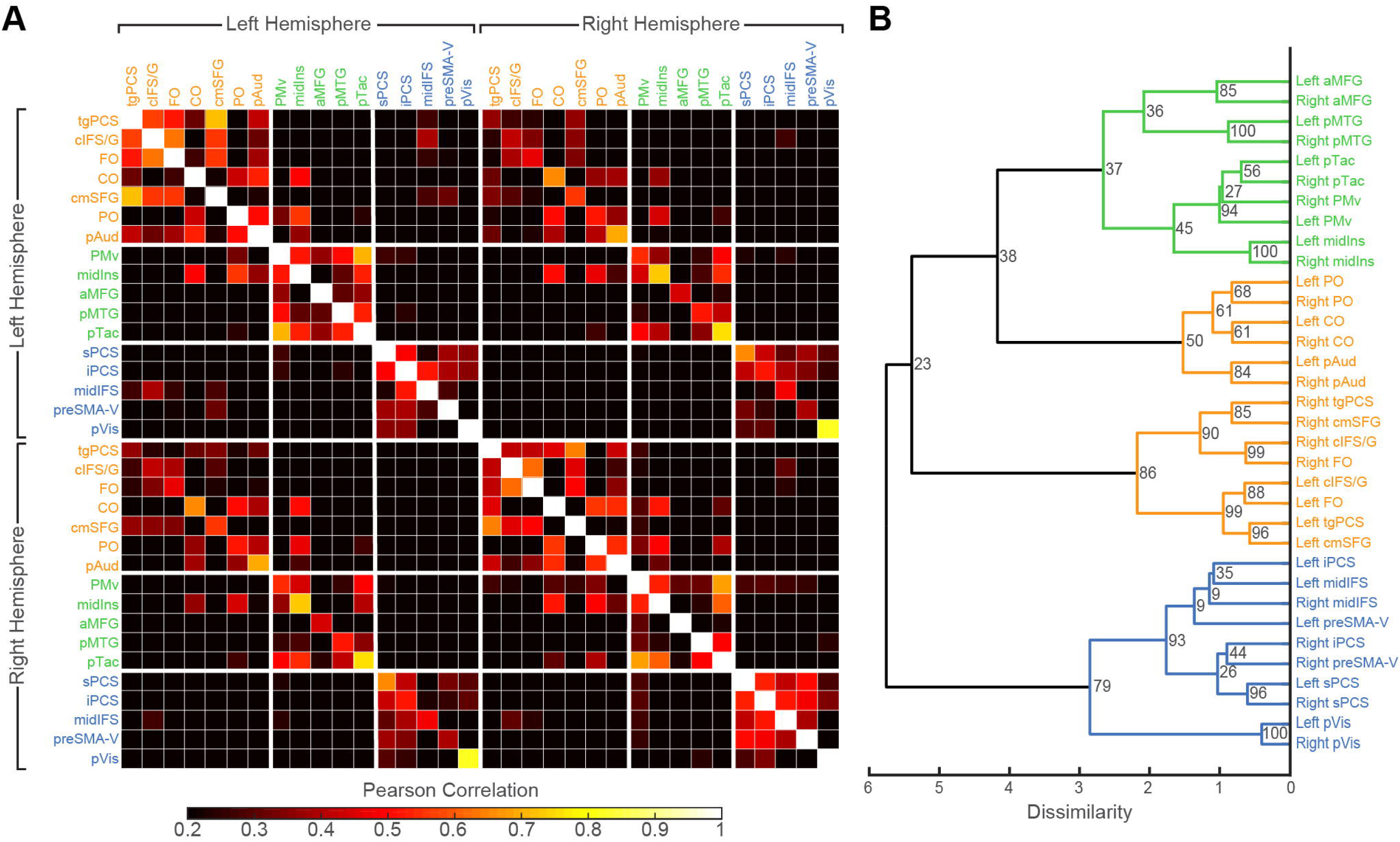
Sensory-biased Functional Connectivity Network Analysis. (A) Adjacency matrix displaying Pearson correlation values between all members of the task-defined modality-biased ROI set. Matrix is thresholded at *r* > 0.2 to remove low and likely non-significant correlation values for visualization only. Thick white lines indicate groups of different modalities. (B) Dendrogram displaying the results of the agglomerative hierarchical cluster analysis (HCA). Input correlation values for the HCA analysis were not thresholded prior to analysis. Numbers at each branch point indicate the proportion of 1000 bootstrap iterations that resulted in the same underlying branch structure. Region names are color-coded according to the modality biased disjunction map they were originally drawn on.

Agglomerative HCA revealed the presence of separate tactile-biased and visual-biased networks and two auditory-biased sub-networks (Figure 6B). In agreement with previously published work (Michalka et al., 2015; Noyce et al., 2017), auditory-biased and visual-biased ROIs were segregated from each other by modality. However, auditory-biased ROIs were segregated into two different subnetworks. Bilateral tgPCS and cIFS/G, regions previously shown to cluster together (Michalka et al., 2015), additionally clustered with bilateral FO and cmSFG. Bilateral pAud, CO and PO formed their own network that was closer in Euclidian distance to the tactile-biased network than the visual-biased or alternate auditory-biased networks; this appears to primarily reflect functional connectivity between the tactile-biased region midIns and each of two auditory-biased regions, CO and PO (see Figure 6A).

The addition of tactile-biased ROIs to the HCA analysis revealed for the first time that a third modality-biased functional network exists alongside the auditory- and visual-biased networks. All tactile ROIs were segregated into a single cluster of tactile-biased network connections. We note that the auditory-biased subnetwork regions with closer linkage to the tactile-biased network lie along the Sylvian fissure and middle-to-posterior insula. Previous rs-fMRI-based research found that these regions cluster into large and stable network nodes belonging to the same ‘motor-auditory’ network (Power et al., 2011; Yeo et al., 2011).

### Group-level Analysis of Sensory-biased WM Recruitment

To illustrate the utility of individual subject-level analyses in revealing sensory-biased WM regions in frontal cortex, group-level analyses were also performed using the same data set. Since prior studies of multiple modalities typically only include two sensory modalities, we present three different pairwise comparisons. Results from the contrasts of auditory > visual, tactile > auditory and tactile > visual are displayed in Figure 7. Group-level analysis of auditory active > visual active reveals no evidence for frontal sensory-biased regions (Figure 7A), even though individual-subject analysis of the same data reveal complex intertwined sensory-biased networks. Posterior pAud and pVis regions survived group-level analysis due to their large size, while auditory-biased CO and PO also survived statistical correction. Tactile active > auditory active again revealed regions in strongly sensory-biased cortex here referred to as pTac and pAud (Figure 7B). Additional group-level tactile recruitment was noted near PMd and pre-supplementary motor area. It is worth noting that PMd and preSMA-T ROIs were drawn in the individual subject analysis but were subsequently eliminated following analysis of WM recruitment – suggesting that these regions are strongly tactile-biased but represent motor response or motor planning processes rather than working memory processes. Auditory-biased recruitment of parietal and lateral occipital cortex was observed; however, this was the result of greater antagonistic deactivation in the tactile condition. Finally, Figure 7C showed a similar pattern to panels A and B – namely that strongly biased pTac and pVis regions are observed but it fails to reveal evidence for sensory-biased regions elsewhere on the cortex surface. These analyses reveal that individual-subject analyses are highly effective in revealing the fine-scale functional architecture of frontal cortex, in conditions for which group-level analyses of the same data fail to reveal functional organization.

**Figure 7:**
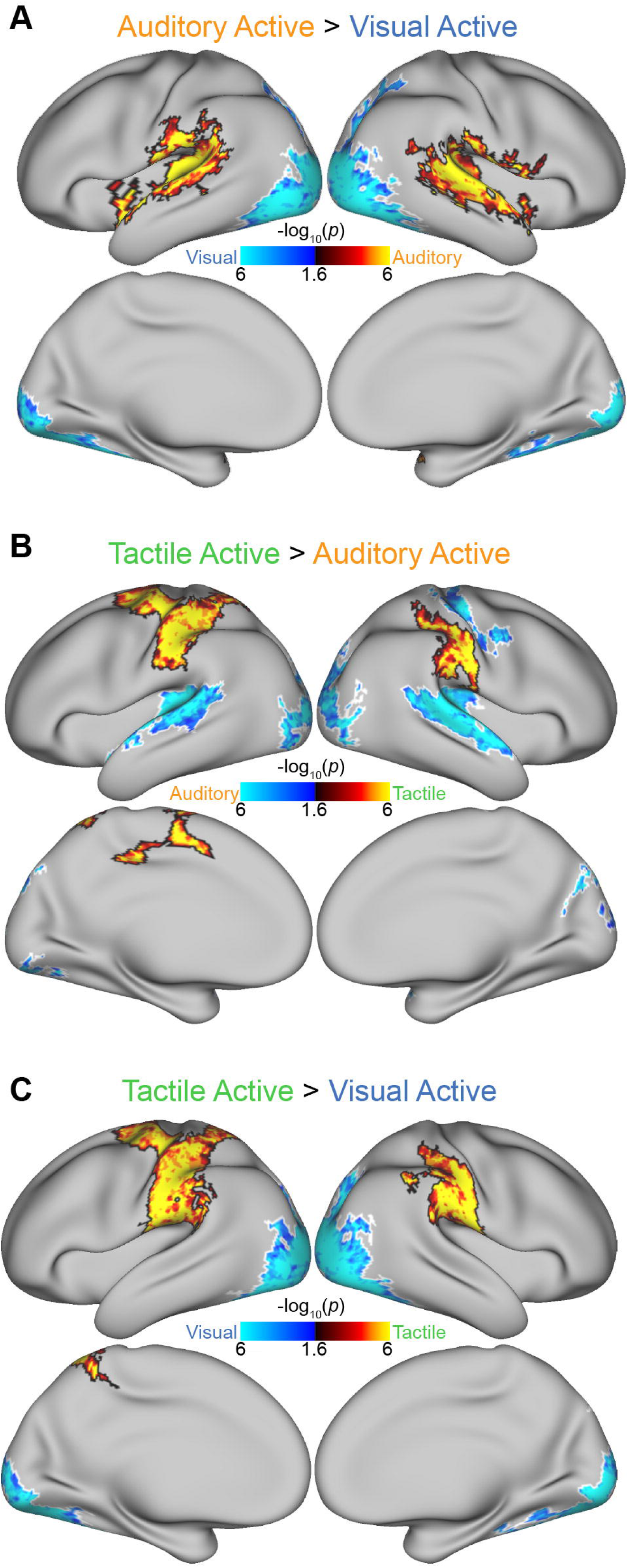
Group-level analysis of Two-modality WM comparisons fail to reveal modality-biased frontal lobe structures that are observed with individual-subject analyses of the same data (Figs 2-4). (A) Auditory active vs. visual active, (B) tactile active vs. visual active and (C) tactile active vs. visual active group analyses were conducted with FSL’s PALM toolset, with additional statistical correction for the analysis of both hemispheres.

### Supramodal Region of Interest Analysis

To examine supramodal WM recruitment across the brain, we performed a vertex-level analysis across the entire cerebral cortex. We examined within-modality sensory WM recruitment (modality A WM vs. modality A sensorimotor control) for each of the three sensory modalities. The z-scores for each modality WM recruitment contrast were combined into a 3-dimensional coordinate system and the resulting data were projected onto the unit sphere (Figure 8A). Vertices that were robustly recruited in all 3 sensory WM modalities projected to a single octant. Across subjects, projecting these supramodal WM vertices from (Figure 8A) back to individual subject surfaces revealed seven reliable cortical regions significantly recruited for all three modality-biased WM conditions. It is important to note that this analysis could identify a cortical vertex as supramodal, even if the earlier (sensory-biased) analyses identified a vertex as sensory-biased; the two definitions are not mutually exclusive. It should also be noted that supramodal or multisensory perceptual regions without robust WM responses would not be captured by any of the ROI definitions. Within individual subjects, three lateral frontal supramodal WM regions were found to partially overlap with visual-biased regions and were thus named by adding -sm (for supramodal) to the visual-biased region name. Three example subjects are depicted in Figure 8B,C while all subjects are depicted in Figures S4 (lateral and posterior views) and S5 (medial view). These regions were: supramodal superior precentral sulcus (sPCS-sm), supramodal iPCS (iPCS-sm) and supramodal midIFS / middle frontal gyrus (midIFS-sm). Anterior and posterior sub-divisions of intraparietal sulcus (aIPS-sm and pIPS-sm, respectively) were reliably found in parietal cortex. Medially, a supramodal region, presupplementary motor area (preSMA-sm) was found amongst the auditory- and visual-biased regions within the general vicinity of preSMA and the medial extent of superior frontal gyrus. In addition to these regions, a supramodal region in anterior insula (aIns-sm) was found. Only aIns-sm does not lie near visual-biased regions identified earlier. In total, seven supramodal WM regions were identified. Their average sizes and the average MNI coordinates of their centroids are reported in Table 1.

**Figure 8:**
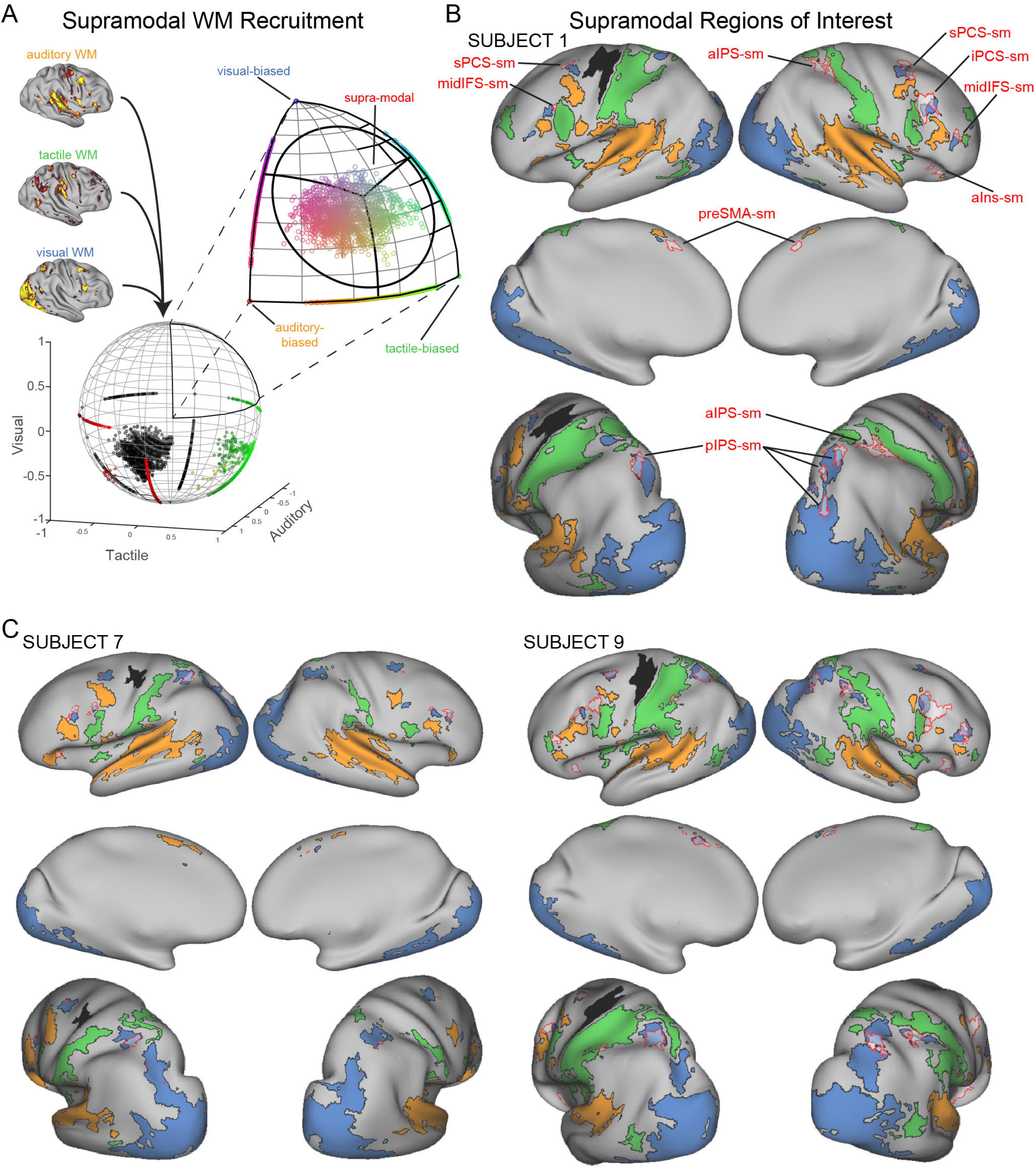
Supramodal WM ROI creation. (A) Schematic of supramodal vertex-wise analysis. Significant *z*-scores (*p* < 0.05) from individual auditory WM > auditory control, tactile WM > tactile control and visual WM > visual control combined into a 3-D coordinate system and were projected to the unit sphere. Vertices are embedded in RGB color space by multiplying their 3D Cartesian coordinate by the matrix [255 0 0; 0 255 0; 0 0 255] to provide a visual representation of uni-sensory, bi-sensory and/or supramodal bias. Note that vertices with 0 or negative *z*-scores in all three modalities are colored black. Points within the circle indicated on the zoomed portion of the sphere are considered supramodal. The ideal supramodal point is indicated by the intersection of the three black lines. (B,C) Supramodal WM ROIs for 3 example subjects are displayed with red borders. Interior white shading is transparent to visualize any overlap between underlying sensory-biased ROIs.

Figure 9 shows the vertex-wise probabilistic overlap map for supramodal WM vertices, with the probabilistic maps for each of the three sensory-biased regions overlaid. On visual inspection, supramodal WM vertices were observed to spare nearly all auditory-biased and tactile-biased ROIs. In contrast to auditory- and tactile-biased ROIs, visual-biased ROIs strongly overlap with supramodal WM vertices. Three multisensory regions in lateral frontal cortex partially overlapped and extended anterior to visual-biased sPCS, iPCS, and midIFS. The supramodal midIFS region also extends into neighboring middle frontal gyrus (MFG). Multisensory recruitment was also observed within Anterior Insula. On the medial surface, preSMA-V was observed to highly overlap with supramodal WM vertices. In the parietal lobe, pVis, a much larger ROI compared to the other visual-biased ROIs, contained a peak of high overlap of supramodal WM vertices in posterior IPS. Additionally, a second parietal supramodal WM region – likely aIPS – appears in the space between pVis and pTac.

**Figure 9:**
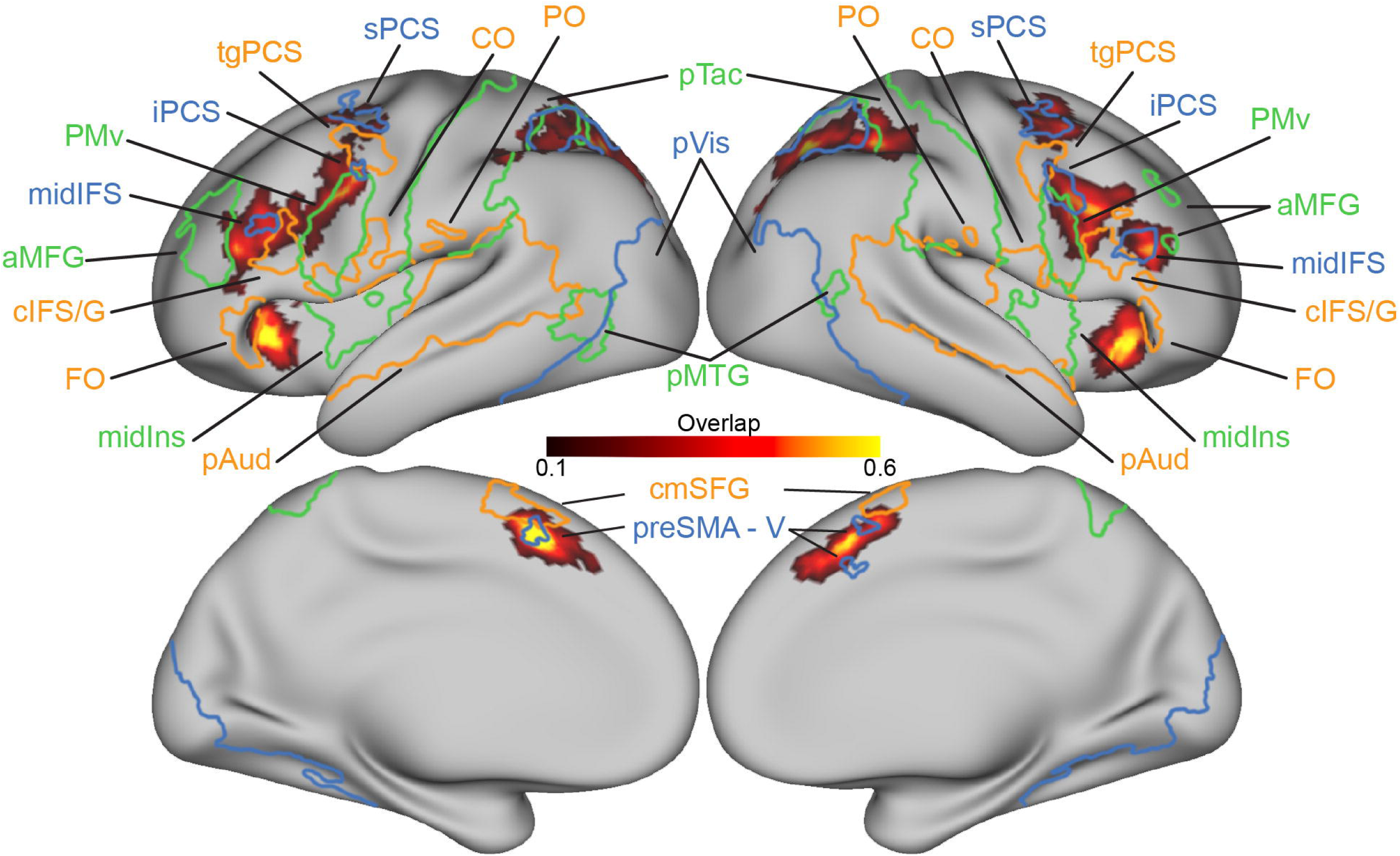
Probabilistic Supramodal Map and Probabilistic Sensory-biased ROIs. The probabilistic overlap map for supramodal ROIs is displayed for the range of 1 (black) to 6 subject overlap (yellow). Auditory-, visual- and tactile-biased ROIs created by thresholded at a 2 subject overlap are displayed in corresponding orange, blue and green borders.

The apparent overlap between supramodal regions of the cortex and visual-biased regions in individual subjects was analyzed in 3 ways. First, the proportion of vertices in each supramodal ROI that was also identified as sensory-biased (via the disjunction analysis) was quantified for each subject, normalized by the area of their supramodal ROI and averaged (Figure 10A). Repeated measures ANOVAs revealed no effect for hemisphere, so they were combined for each ROI. Note that aIns-sm contains very few sensory-biased vertices and thus is distinctly supramodal. Similarly, 75% of vertices in preSMA-sm did not exhibit a sensory bias. Approximately 50% of vertices in frontal supramodal regions midIFS-sm, sPCS-sm, iPCS-sm, and parietal aIPS-sm did not exhibit a sensory-bias. Nevertheless, except for aIns-sm, at least 20% of vertices within each supramodal WM ROI were also identified as visual-biased. However, this proportion only reached significance for pIPS-sm (Table 2). A two*-*way ANOVA for ROI and modality showed a significant main effect for modality (*F*_(2,180)_ = 55.96, *p* = 1.26×10^−77^). Tukey’s HSD post-hoc analysis revealed that significantly greater proportions of midIFS-sm, sPCS-sm and pIPS-sm were visual-biased when compared to auditory- or tactile-biased. iPCS-sm had significantly more area that was visual-biased but equal proportions of tactile-biased or auditory-biased area. aIns-sm, preSMA-sm and aIPS-sm showed no significant differences for modality. Note that aIns-sm contains only auditory-biased overlap, preSMA-sm contains only auditory- and visual-biased overlap and aIPS-sm contains only visual- and tactile-biased overlap.

**Figure 10:**
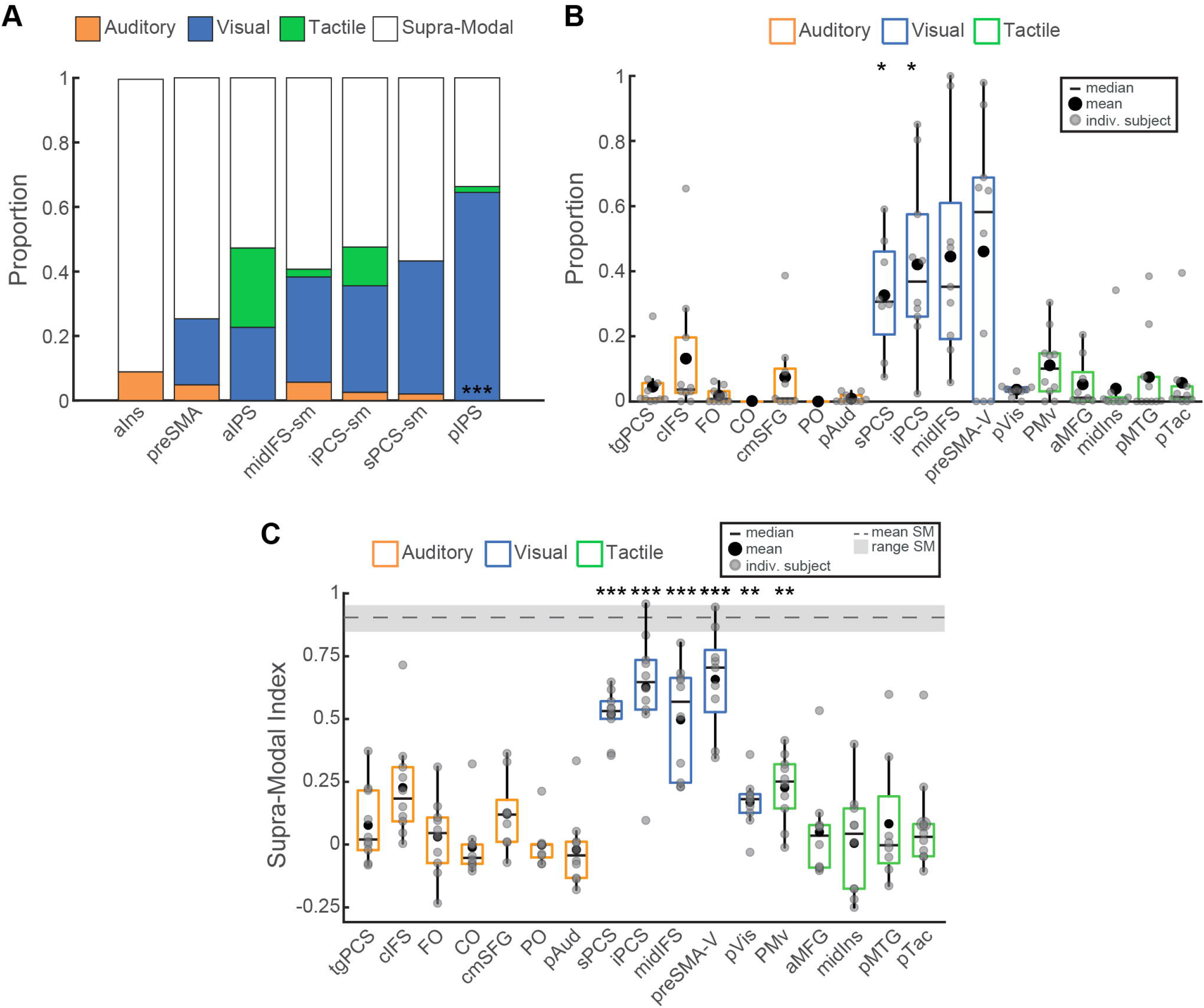
Supramodal and Sensory-biased Colocalization Metrics. (A) The proportion of each supramodal ROI that was also identified as sensory-biased (any modality) is depicted by stack bar plots. Each ROIs proportions were normalized to 1 by each subject’s supramodal area prior to aggregating. (B) The proportion of each sensory-biased ROI that was also identified as supramodal is depicted by box plots. Each ROIs proportions were normalized by each subject’s area prior to aggregating. (C) SMI values calculated for each ROI. To provide context, the average (dashed line) and range (gray shading) of SMI values across all supramodal ROIs is included. *p < 0.05, **p < 0.01, ***p < 0.001, Student’s *t*-tests, Holm-Bonferroni corrected.

**Table 2:** Proportion of Supramodal ROIs Overlapping with Sensory-biased ROIs. The definition for supramodal WM is not exclusive with the definitions of sensory-biased WM. The proportion of each supramodal WM ROI that overlapped sensory-biased WM ROIs in each sensory modality was analyzed at the individual-subject level and summarized here.

| ROI | Repeated Measures ANOVA<br>(main effect of hemisphere) | Average Proportion ± SD<br>(Student's *t*-test) | | |
| --- | --- | --- | --- | --- |
|  |  | Aud | Vis | Tac |
| Left alns-sm | $F_{(1,21)} = 2.18$;<br>$p = 0.16$ | $0.09 \pm 0.23$<br>$t(9) = 1.25$;<br>$p = 1$ | 0 | $0.004 \pm 0.02$<br>$t(9) = 1$;<br>$p = 1$ |
| Right alns-sm |  |  |  |  |
| Left preSMA-sm | $F_{(1,24)} = 1.68$;<br>$p = 0.21$ | $0.05 \pm 0.08$<br>$t(9) = 1.91$;<br>$p = 0.89$ | $0.21 \pm 0.22$<br>$t(9) = 2.94$;<br>$p = 0.23$ | 0 |
| Right preSMA-sm |  |  |  |  |
| Left aIPS-sm | $F_{(1,21)} = 2.08$;<br>$p = 0.16$ | 0 | $0.23 \pm 0.3$<br>$t(8) = 2.26$;<br>$p = 0.59$ | $0.25 \pm 0.21$<br>$t(8) = 3.53$;<br>$p = 0.13$ |
| Right aIPS-sm |  |  |  |  |
| Left midIFS-sm | $F_{(1,21)} = 0.14$;<br>$p = 0.71$ | $0.6 \pm 0.06$<br>$t(9) = 3.12$;<br>$p = 0.18$ | $0.33 \pm 0.27$<br>$t(9) = 3.89$;<br>$p = 0.07$ | $0.2 \pm 0.03$<br>$t(9) = 2.75$;<br>$p = 0.29$ |
| Right midIFS-sm |  |  |  |  |
| Left iPCS-sm | $F_{(1,27)} = 0.64$;<br>$p = 0.43$ | $0.2 \pm 0.03$<br>$t(9) = 2.48$;<br>$p = 0.42$ | $0.33 \pm 0.3$<br>$t(9) = 3.54$;<br>$p = 0.12$ | $0.12 \pm 0.11$<br>$t(9) = 3.33$;<br>$p = 0.14$ |
| Right iPCS-sm |  |  |  |  |
| Left sPCS-sm | $F_{(1,18)} = 1 \times 10^{-4}$;<br>$p = 0.99$ | $0.02 \pm 0.04$<br>$t(8) = 1.43$;<br>$p = 1$ | $0.41 \pm 0.34$<br>$t(8) = 3.64$;<br>$p = 0.12$ | 0 |
| Right sPCS-sm |  |  |  |  |
| Left pIPS-sm | $F_{(1,24)} = 1.01$;<br>$p = 0.32$ | 0 | **$0.64 \pm .2$**<br>**$t(8) = 9.62$;**<br>**$p = 0.0002$** | $0.02 \pm 0.03$<br>$t(8) = 1.88$;<br>$p = 0.89$ |
| Right pIPS-sm |  |  |  |  |

A second analysis performed proportion analysis from the opposite perspective by examining the proportion of vertices in each sensory-biased ROI that was identified as supramodal WM (Figure 10B). A repeated measures ANOVA revealed no effect of hemisphere, so all ROIs were combined across hemispheres. A two-way ANOVA revealed a significant effect for ROI (*F*_(16,163)_ = 9.98, *p* = 1.23×10^−16^). Subsequent Tukey’s HSD post-hoc tests revealed that significantly greater proportions of iPCS, midIFS and preSMA-V were supramodal compared to all other regions. However, variability within ROIs was such that Student’s *t-*tests revealed only sPCS, iPCS and pVis were significantly different from 0 (Table 3).

**Table 3:** Statistical analysis of supramodal proportion within sensory-biased ROIs. The proportion of each sensory-biased WM ROI that overlapped with supramodal WM ROIs was analyzed at the individual-subject level and summarized here.

| ROI | Repeated Measures ANOVA<br>(main effect of hemisphere) | Proportion<br>(mean ± SD) | Student's *t*-test |
| --- | --- | --- | --- |
| Left tgPCS | $F_{(1,9)} = 1.76; p = 0.23$ | $0.05 \pm 0.08$ | $t_{(9)} = 1.8; p = 0.74$ |
| Right tgPCS |  |  |  |
| Left cIFS/G | $F_{(1,8)} = 4.3; p = 0.07$ | $0.13 \pm 0.21$ | $t_{(9)} = 2.02; p = 0.63$ |
| Right cIFS/G |  |  |  |
| Left FO | $F_{(1,9)} = 1.7; p = 0.23$ | $0.02 \pm 0.02$ | $t_{(9)} = 2.44; p = 0.41$ |
| Right FO |  |  |  |
| Left pAud | $F_{(1,9)} = 0.24; p = 0.63$ | $0.009 \pm 0.01$ | $t_{(8)} = 2.13; p = 0.62$ |
| Right pAud |  |  |  |
| Left CO | $F_{(1,9)} = 2.02; p = 0.19$ | $0.001 \pm 0.003$ | $t_{(9)} = 1.42; p = 0.67$ |
| Right CO |  |  |  |
| Left cmSFG | $F_{(1,6)} = 0.8; p = 0.41$ | $0.08 \pm 0.13$ | $t_{(8)} = 1.79; p = 0.64$ |
| Right cmSFG |  |  |  |
| Left PO | No Overlap | 0 | No Overlap |
| Right PO |  |  |  |
| Left sPCS | $F_{(1,7)} = 0.17; p = 0.69$ | $0.33 \pm 0.18$ | **$t_{(7)} = 5.25; p = 0.02$** |
| Right sPCS |  |  |  |
| Left iPCS | $F_{(1,9)} = 0.11; p = 0.75$ | $0.42 \pm 0.26$ | **$t_{(9)} = 5.13; p = 0.01$** |
| Right iPCS |  |  |  |
| Left midIFS | $F_{(1,6)} = 0.02; p = 0.9$ | $0.46 \pm 0.34$ | $t_{(8)} = 3.97; p = 0.06$ |
| Right midIFS |  |  |  |
| Left pVis | $F_{(1,9)} = 1.89; p = 0.2$ | $0.07 \pm 0.02$ | **$t_{(9)} = 4.7; p = 0.01$** |
| Right pVis |  |  |  |
| Left preSMA-V | $F_{(1,7)} = 0.22; p = 0.65$ | $0.46 \pm 0.338$ | $t_{(9)} = 3.83; p = 0.06$ |
| Right preSMA-V |  |  |  |
| Left PMv | $F_{(1,9)} = 0.03; p = 0.88$ | $0.11 \pm 0.1$ | $t_{(9)} = 3.43; p = 0.09$ |
| Right PMv |  |  |  |
| Left midIns | $F_{(1,9)} = 2.37; p = 0.07$ | $0.04 \pm 0.11$ | $t_{(9)} = 1.16; p = 0.57$ |
| Right midIns |  |  |  |
| Left aMFG | $F_{(1,5)} = 0.21; p = 0.67$ | $0.08 \pm 0.18$ | $t_{(8)} = 2.1; p = 0.63$ |
| Right aMFG |  |  |  |
| Left pMTG | $F_{(1,9)} = 1.08; p = 0.33$ | $0.07 \pm 0.12$ | $t_{(9)} = 1.79; p = 0.74$ |
| Right pMTG |  |  |  |
| Left pTac | $F_{(1,9)} = 0.78; p = 0.40$ | $0.06 \pm 0.12$ | $t_{(9)} = 1.49; p = 0.67$ |
| Right pTac |  |  |  |

A third analysis examined the SMI values across every vertex in each sensory-biased ROI for each subject. No main effect for hemisphere was found for SMI values across ROIs. A main effect for ROI was found (*F*_(16,164)_ = 17.1, *p* = 3.99×10^−26^) and Tukey’s post-hoc tests indicated that sPCS, iPCS, midIFS and preSMA-V each possessed significantly higher average SMI than all other ROIs. Student’s *t-*tests showed that no auditory-biased ROI possessed significant supramodal characteristics (Figure 10C, Table 4). Only a single tactile-biased ROI, left PMv, was found to possess significant SMI. However, all visual-biased ROIs showed significantly high SMI values. Collectively, these results demonstrate that supramodal WM vertices in frontal cortex tend to also be visual-biased (Figure 10a) and that visual-biased vertices in frontal cortex tend to exhibit supramodal WM properties (Figure 10b,c); supramodal processing does not exhibit this relationship with the auditory or tactile modalities.

**Table 4:**
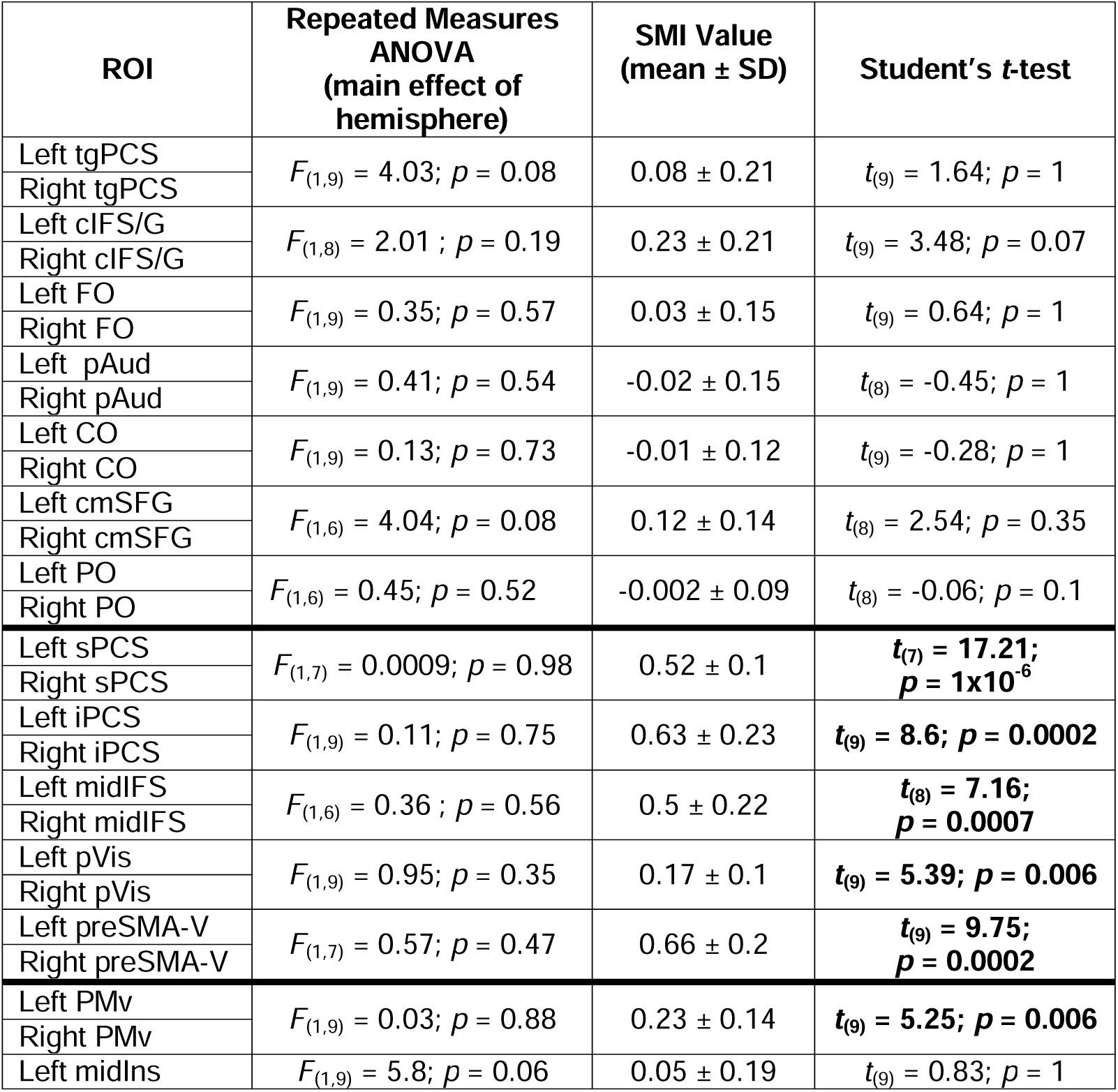
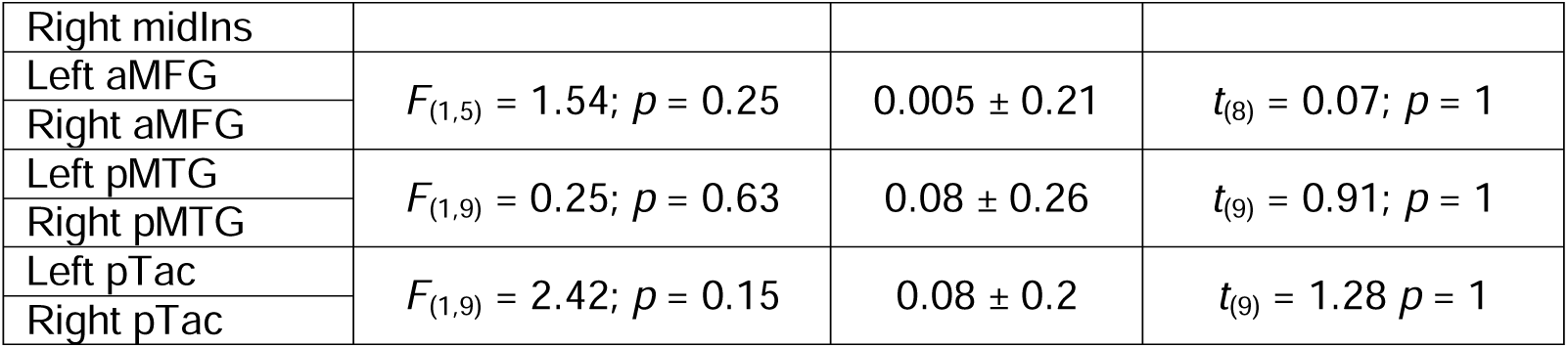
Statistical analysis of SMI values by sensory-biased ROI. For each subject and each sensory-biased ROI, a supramodal WM index (SMI) was computed from the t-stats of the active vs. sensorimotor control contrast in each sensory modality (See Materials and Methods). Subject-level measures were analyzed with a t-test to determine the significance of the SMI values (vs. 0) for each ROI.

The cortical architecture of sensory-biased working memory and multiple sensory modality working memory regions are summarized in the schematic of Figure 11.

**Figure 11:**
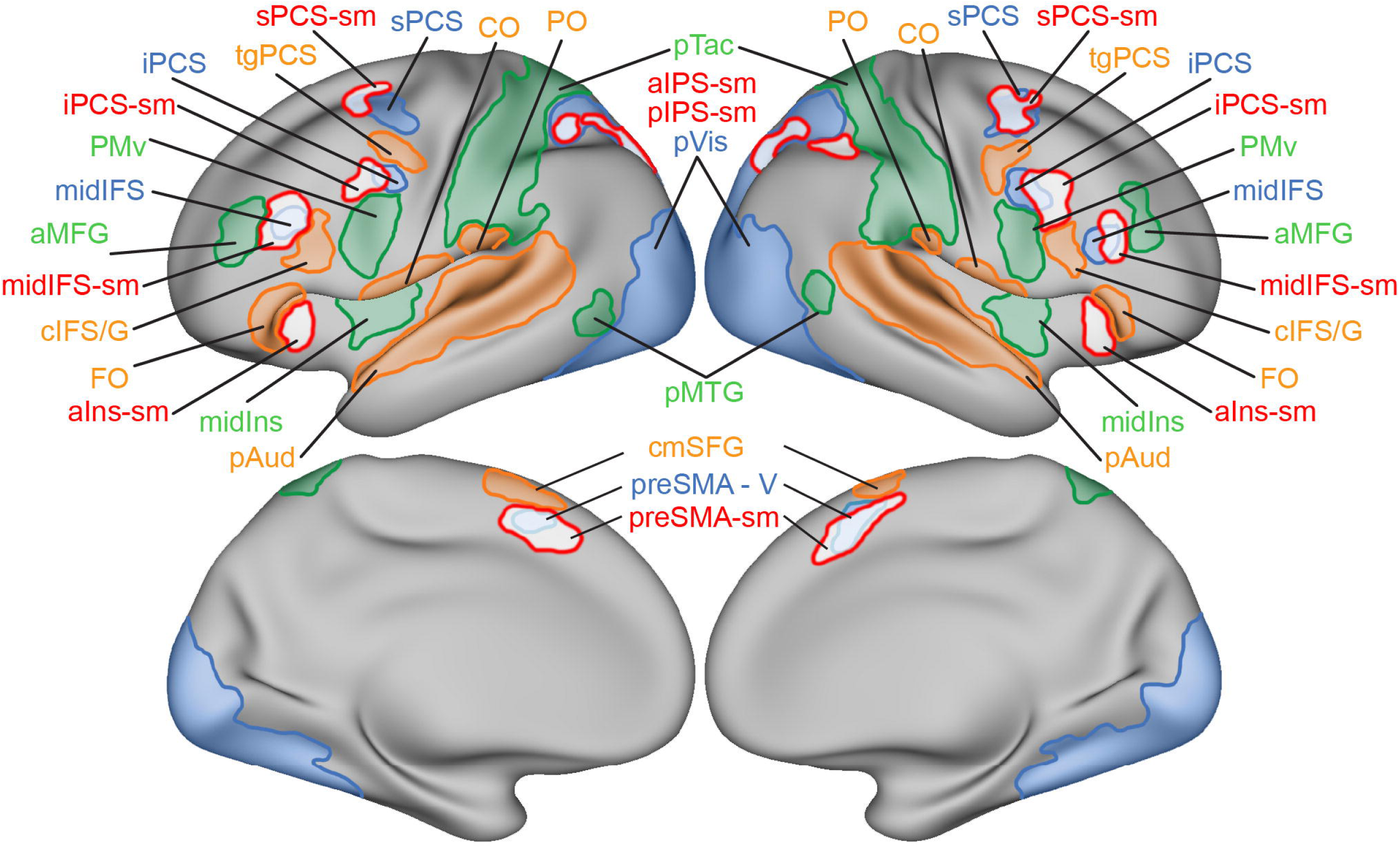
Schematic Representation of Sensory-biased and Supramodal ROIs. All ROIs are depicted in their approximate location and size to serve as a schematic representation of the topography of sensory-biased and supramodal WM regions across the cortex. Supramodal region (red border) are shown with a semi-transparent white fill to allow the overlapping visual regions to be seen. Overlap between visual and supramodal ROIs was drawn jointly according to individual subject and probabilistic labels.

## Discussion

We examined sensory working memory processing in the visual, auditory, and tactile modalities. Our analyses reveal multiple, bilateral frontal cortical structures that are biased for sensory modality in WM tasks. We also observe a broad network of seven supramodal WM structures that are recruited for working memory in all three sensory modalities. Within-subject analyses were critical as the sensory-biased frontal regions are relatively small, are interleaved between different modalities, and exhibit anatomical heterogeneity across individuals. Group-averaged analyses of the same data set blur out frontal lobe distinctions for sensory modality, leaving sensory-biased structures only in posterior cortices. Nevertheless, examination of individual brains reveals a consistent functional organization of sensory-biased and supramodal WM regions (Figure 11; Table 1). The observed visual-biased and auditory-biased structures replicate our prior report (Noyce et al., 2022; see also Michalka et al., 2015; Noyce et al., 2017), but here we also report the existence of tactile-biased structures located adjacent to the visual- and auditory-biased frontal lobe regions. Our functional definitions allow for a voxel or ROI to be categorized as both sensory-biased and supramodal (responsive to WM in all 3 modalities). Many supramodal WM voxels are also visual-biased (Figure 10a) and many visual-biased voxels in frontal cortex exhibit supramodal WM properties (Figure 10b,c); neither the auditory nor tactile modality exhibit this relationship with supramodal processing. This observation is consistent with prior reports indicating strong links between multiple-demand regions and the visual-biased network (Noyce et al., 2017, 2022; Tobyne et al., 2017).

Five bilateral auditory-biased frontal lobe regions were observed in this study: transverse gyrus intersecting the precentral sulcus (tgPCS), caudal inferior frontal sulcus/gyrus (cIFS/G), frontal operculum (FO), central operculum (CO), and the caudal medial portion of the superior frontal gyrus (cmSFG). Bilateral visual-biased frontal lobe regions were observed in superior precentral sulcus (sPCS), inferior precentral sulcus (iPCS), and the middle of inferior frontal sulcus (mid-IFS). These eight bilateral sensory-biased regions were also reported in our prior study (Noyce et al., 2022) and six of these regions were independently observed (Assem et al., 2022). Here, our analysis revealed a small visual-biased region within the pre-supplementary motor area, that we refer to as preSMA-v. This region was observed in some participants in (Noyce et al., 2022), but was not significant in group-level analysis of individual subjects.

The inclusion of the tactile modality revealed 2 bilateral tactile-biased WM frontal lobe regions –ventral premotor (PMv) and anterior middle frontal gyrus (aMFG), which were localized within previously unassigned ‘blank’ areas of the sensory-biased frontal cortical schematic (Figure 5). We also observed tactile-biased WM regions in mid-Insula (midIns), posterior middle temporal gyrus / lateral occipital cortex (pMTG), as well as large swath of anterior parietal lobe (pTac), which likely includes SI and SII. PMv lies within a region shown to possess differential connectivity with primary motor cortex (Glasser et al., 2016), and while this region is typically described as premotor (e.g., Amunts et al., 2010; Glasser et al., 2016), we found that PMv is significantly recruited for tactile WM (vs. sensory motor control). Previous research found evidence for tactile WM signals in ventrolateral frontal cortex in non-human primates and human populations (Romo et al., 1999; Kostopoulos et al., 2007; Sörös et al., 2007; Wu et al., 2018). Resting-state functional connectivity of sensory-biased regions, revealed that the tactile-biased frontal and posterior regions formed a network, separate from the visual- and auditory-biased networks. To our knowledge, this is the first fMRI report of the WM recruitment of distinct auditory, visual- and tactile-biased regions in human frontal lobes.

The tactile-biased regions PMv, aMFG, and midIns are larger than frontal visual-biased and auditory-biased regions; this is prominent in left hemisphere, suggesting a strong contralateral drive since tactile stimuli were presented only to the right index finger. Additionally, we employed a haptic exploration paradigm, which might evoke more extensive networks than passive WM paradigms. It will be critical for future WM studies comparing sensory modalities to include a broader range of tactile stimuli and paradigms.

We observed supramodal WM activation in five bilateral frontal cortical areas – anterior insula (antIns), pre-supplementary motor area (pre-SMA), middle inferior frontal gyrus / middle frontal gyrus (midIFS/MFG), superior pre-central sulcus (sPCS), and inferior pre-central sulcus (iPCS) – and two parietal regions – anterior intraparietal sulcus (aIPS) and posterior intraparietal sulcus (pIPS). Our experiments presented for only one sensory modality at a time and thus we could not examine multi-sensory integration (e.g., (Stein & Meredith, 1993; Wallace et al., 1996; Beauchamp, 2005). Our experimental design also does not permit us to operationally distinguish between supramodal WM and multi-demand processing (Duncan, 2010); additional experimental approaches (e.g., Goebel & van Atteveldt, 2009) would be needed. We conjecture that supramodal WM regions comprise a subset of the multiple-demand network; our seven supramodal WM regions roughly align with seven of the ten regions defined as the ‘core’ multiple-demand network (Assem et al., 2020, 2022).

We examined the anatomical relationship between supramodal regions and sensory-biased regions. One prior study reported that visual-biased and auditory-biased frontal regions lie posterior to multiple-demand regions (Assem et al., 2022); in contrast, we previously reported that visual-biased (but not auditory-biased) frontal lobe regions exhibited multiple demand properties (Noyce et al., 2017, 2022). Here, we find evidence supporting both prior reports – supramodal WM / multiple-demand frontal lobe regions are centered somewhat anterior to the sensory-biased regions, but the visual-biased and supramodal regions also exhibit significant overlap. This supramodal overlap does not broadly extend to the auditory- or tactile-biased regions. Specifically, the individual-subject analysis demonstrates that the visual-biased frontal ROIs had roughly 40% of their vertices identified as supramodal WM, while none of the auditory-biased or tactile-biased ROIs had more than 20% of their vertices identified as supramodal WM. The extent of visual-biased frontal lobe ROIs appears somewhat smaller than that of auditory-biased and tactile-biased frontal lobe regions (Table 1), so the overlap between visual-biased and supramodal representations does not simply reflect more extensive frontal lobe recruitment in the visual task than in the other two modalities.

Comparison of the probabilistic ROIs for visual-biased and supramodal WM responses shows that the centroids of the regions are noticeably shifted. Visual-biased sPCS, iPCS, and midIFS each appear centered posterior with respect to the nearest supramodal regions. This suggests a possible posterior-to-anterior functional gradient between visual-biased and supramodal/multiple demand representations in each of these three regions of lateral prefrontal cortex. The two supramodal regions in intraparietal sulcus appear to lie to the lateral side of the visual-biased regions, which may relate to a previously reported medial-to-lateral functional gradient between visuo-spatial attention and WM (Brissenden et al., 2018; Lefco et al., 2020).

Collectively, these results raise the question of why the visual modality is more tightly anatomically integrated with supramodal representations than are the auditory or tactile modalities. Even though this finding is consistent with our visual vs. auditory analysis from two prior datasets (Michalka et al., 2015; Noyce et al., 2017), the observation of an asymmetry between the sensory modalities is counter-intuitive. If our conjecture is correct that supramodal WM regions comprise a subset of the multiple-demand network, this would imply a tight link between visual and multiple-demand processing. In prior resting-state functional connectivity analyses, we have reported that nodes of the visual-biased network exhibit robust connectivity with purported domain-general brain regions, while nodes of the auditory-biased network exhibit robust connectivity with purported language network regions (Tobyne et al., 2017; Noyce et al., 2022). Perhaps each sensory modality has privileged connectivity with one non-sensory network. We have not examined tactile-biased connectivity more broadly; one could speculate that robust connections may occur with the motor network. Some behavioral studies have reported a tighter link between domain-general cognitive processing and visual working memory than with the auditory modality in adults (Bo et al., 2011) and children (Alhamdan et al., 2023), but other potential behavioral implications of this visual-supramodal network link deserve broader investigation.

These findings demonstrate extended cortical networks that maintain a preference for each of three sensory modalities during WM tasks. The extended WM networks are consistent with the view that WM mechanisms are broadly distributed across the brain (Postle, 2015; Christophel et al., 2017; Brissenden et al., 2021). In a broad sense, the observed preferences for sensory modality WM align with Baddeley’s arguments for multiple content-specific WM stores (Baddeley, 2012), although the current study does not directly isolate phonological or visuospatial WM. It is notable that many of these sensory-biased regions lie anatomically adjacent to supramodal WM/ multiple-demand regions, and the visual-biased regions partially overlap the multiple-demand regions, possibly forming a functional gradient. One important line of future inquiry is to distinguish different aspects of WM, such as which regions reflect the contents of WM in each modality and which processes (e.g., encoding, maintenance and retrieval) are supported by each region. Here, we used a N-back paradigms which can produce robust activation, but don’t easily isolate specific component WM processes. Network recruitment might also vary with different types of stimuli. Also important is to examine how these networks are recruited to support multisensory integration and WM for multisensory objects.

## ACKNOWLEDGEMENTS

This work was generously funded through grants provided by the National Science Foundation (SMA-0835976 to DCS), National Eye Institute (R01-EY022229 to DCS), National Institute of Neurological Disorders and Stroke (F31-NS103306 to SMT), and National Institute on Drug Abuse (T90-DA032484 to SMT). The authors wish to thank Vaibhav Tripathi, David Beeler, and Thomas Possidente for helpful discussions.

Dr. Tobyne’s current affiliation is Linus Health, Boston, MA 02210. Dr. Brissenden’s current affiliation is University of Michigan. Dr. Noyce’s current affiliation is Carnegie Mellon University.

## SUPPLEMENTAL DATA FIGURE LEGENDS

**Figure S1: Working Memory vs. Control Condition Activation in Sensory-Biased ROIs in their preferred modality.** Test of WM activation for each candidate ROI. (A) contrast of visual WM vs. visual control within visual-biased ROIs, (B) auditory WM vs. auditory control within auditory-biased ROIs, and (C) tactile WM vs. tactile control within tactile-biased ROIs, across all subjects. *p < 0.05, **p < 0.01, ***p < 0.001, Student’s *t*-tests, Holm-Bonferroni corrected. CO, PPC, PMd, and preSMA-T are non-significant and thus were excluded from further analysis.

**Figure S2: Full Sensory-biased ROI Sets for all subjects.** Auditory-, visual- and tactile-biased ROIs drawn from their respective maps are displayed for all 10 subjects.

**Figure S3: Major anatomical landmarks for brain renderings.** Sulci are labeled on the left hemisphere, and gyri are labeled on the right hemisphere.

**Figure S4: Sensory-biased WM and Supramodal WM ROI for all subjects (lateral, posterior views).** Multisensory and Auditory-, visual- and tactile-biased ROIs are displayed for all 10 subjects.

**Figure S5: Sensory-biased WM and Supramodal WM ROI for all subjects (medial view).** Multisensory and Auditory-, visual- and tactile-biased ROIs are displayed for all 10 subjects.

