## Supplementary figures and images for "Combined Auditory, Tactile, and Visual fMRI Reveals Sensory-Biased and Supramodal Working Memory Regions in Human Frontal Cortex"

### Supplemental Figure 1

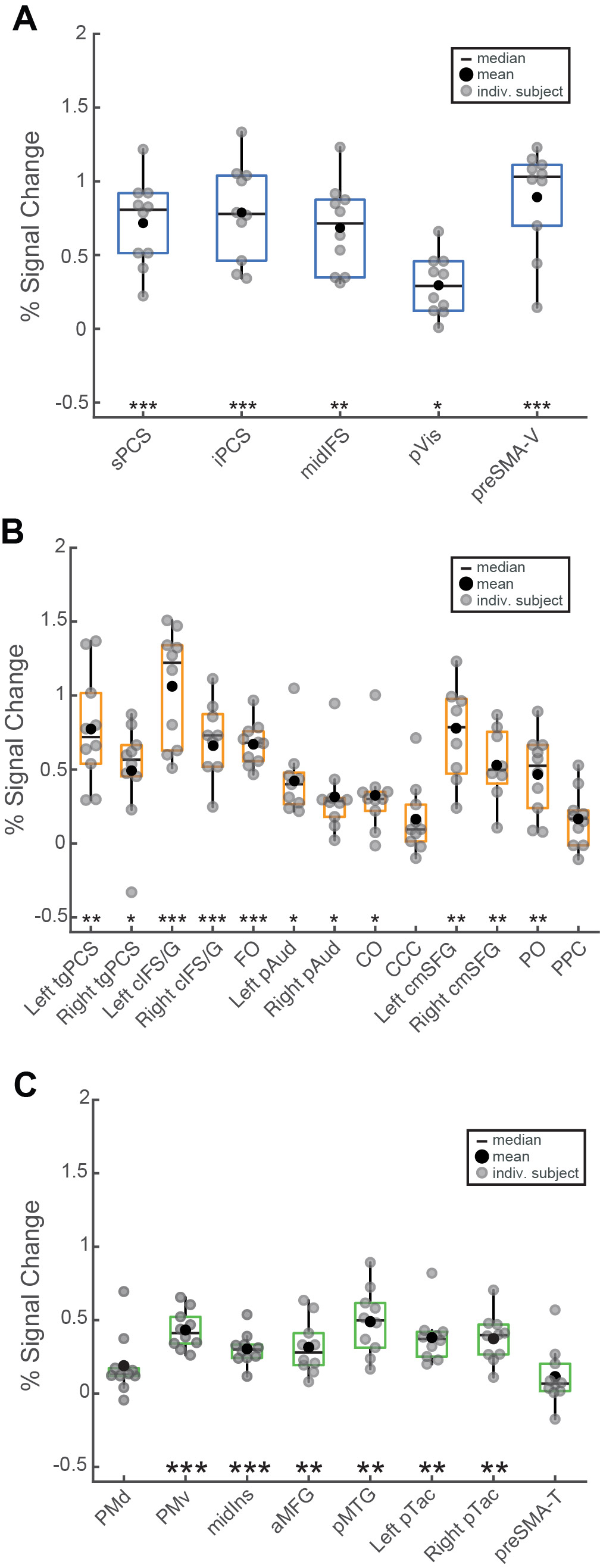

### Supplemental Figure 2

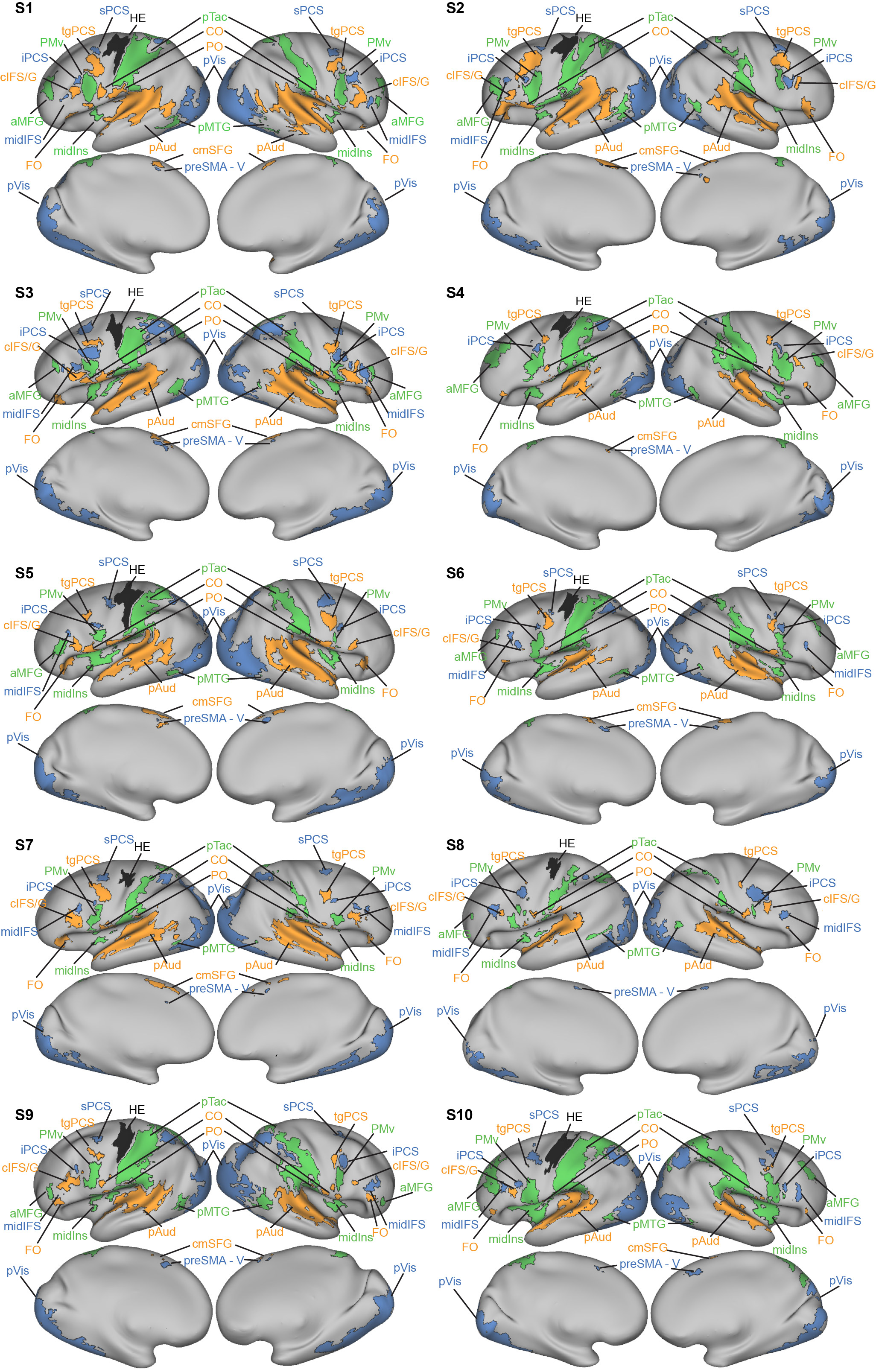

### Supplemental Figure 3

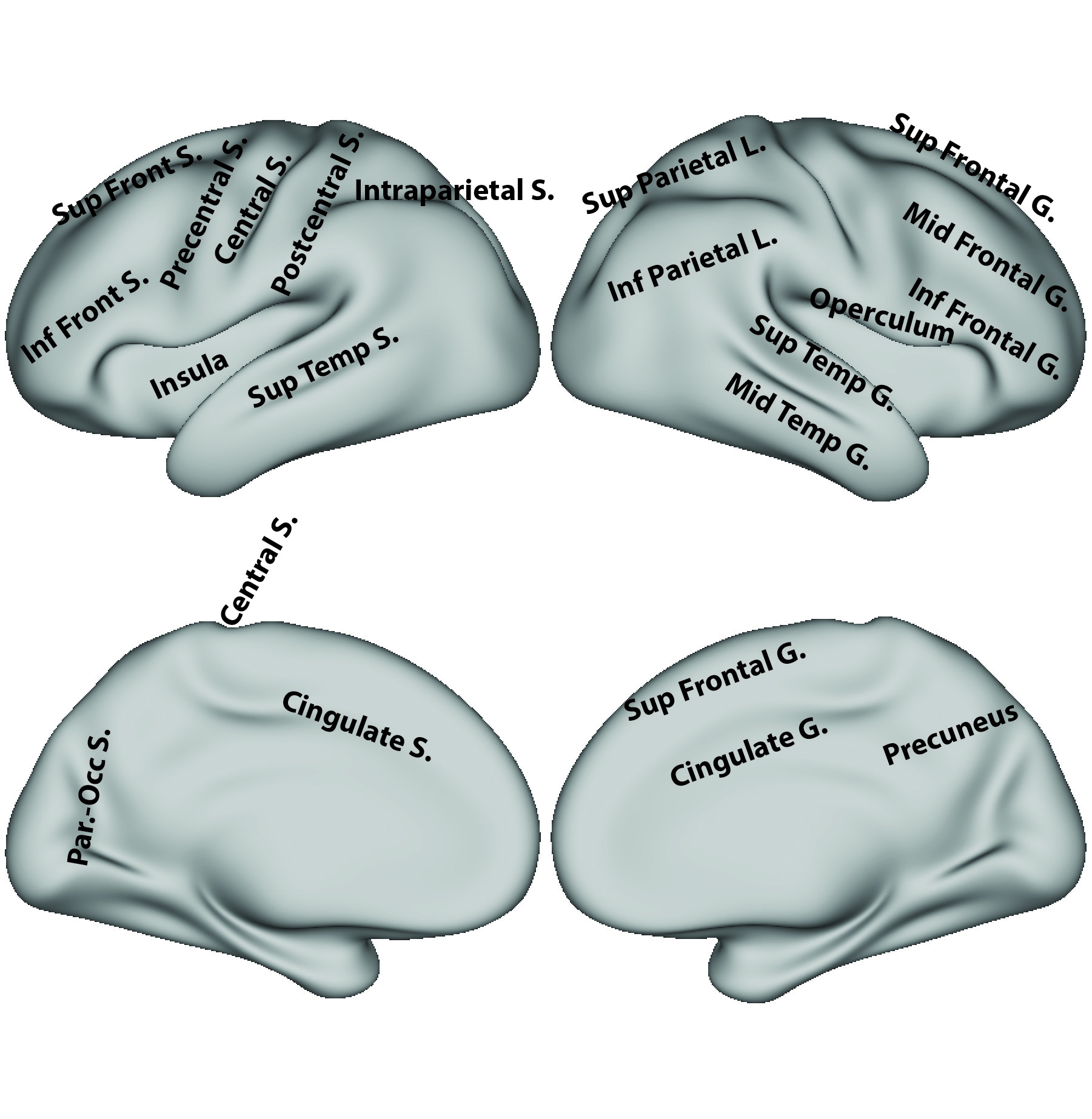

### Supplemental Figure 4

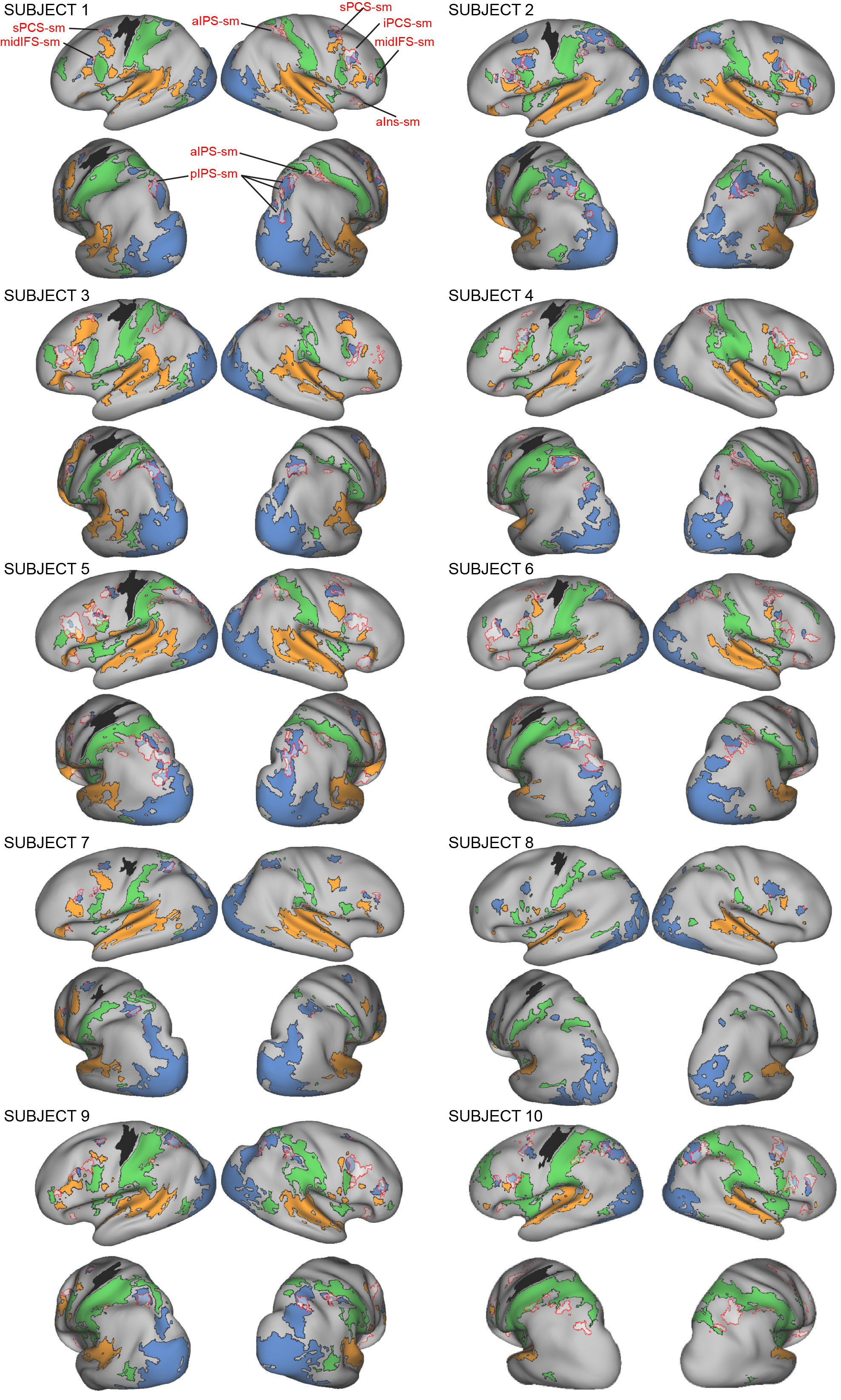

### Supplemental Figure 5

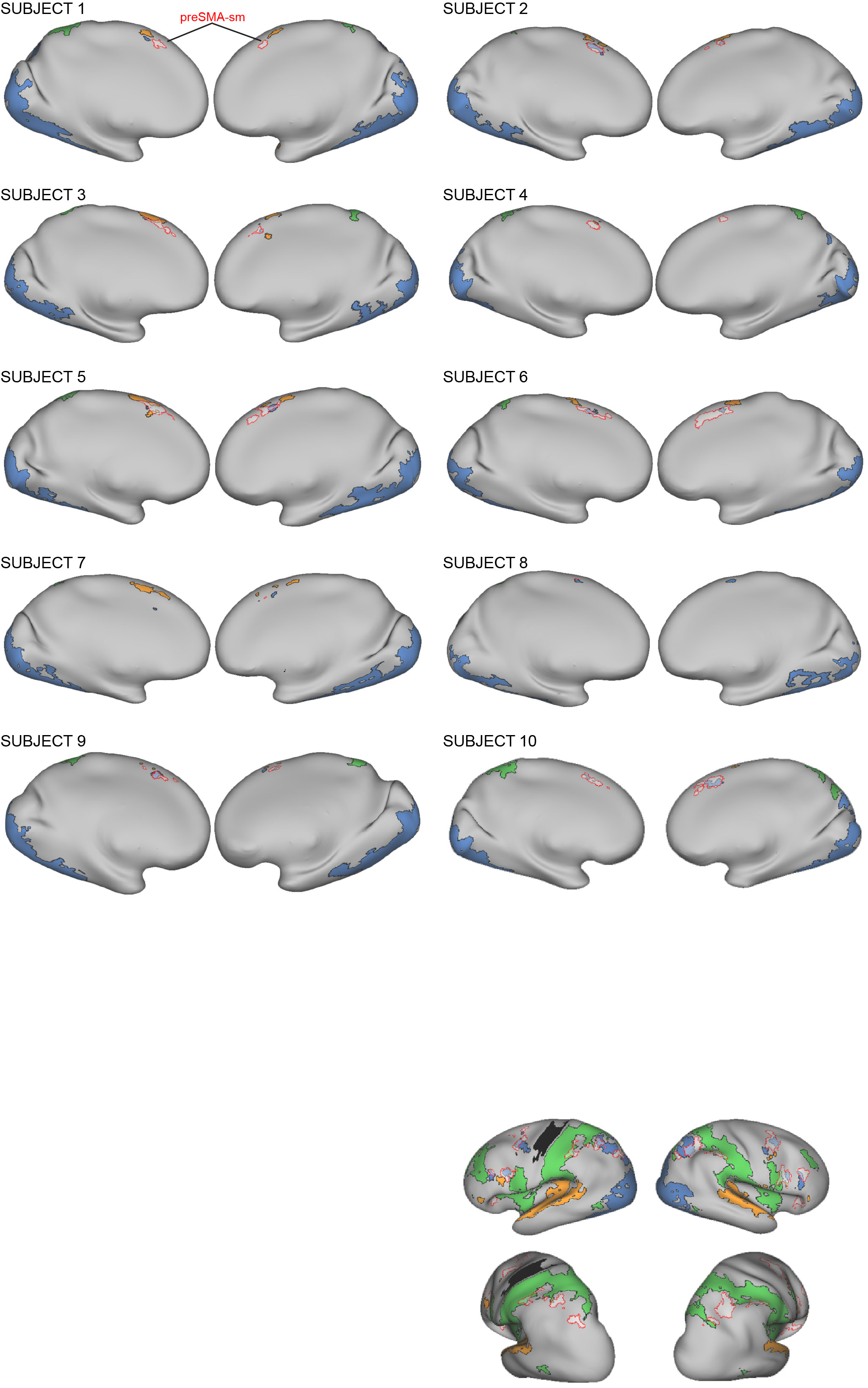
